# The impact of geographic targeting of oral cholera vaccination in sub-Saharan Africa: a modeling study

**DOI:** 10.1101/617076

**Authors:** Elizabeth C. Lee, Andrew S. Azman, Joshua Kaminsky, Sean M. Moore, Heather S. McKay, Justin Lessler

**Author notes:** Correspondence to: Elizabeth C. Lee, Department of Epidemiology, Johns Hopkins Bloomberg School of Public Health, Baltimore, MD, 21205, USA.

## Abstract

**Background:** In May 2018, the World Health Assembly committed to reducing worldwide cholera deaths by 90% by 2030. Oral cholera vaccine (OCV) plays a key role in reducing the near-term risk of cholera, although global supplies are limited. Characterizing the potential impact and cost-effectiveness of mass OCV deployment strategies is critical for setting expectations and developing cholera control plans that maximize chances of success.

**Methods and Findings:** We compared the projected impacts of vaccination campaigns across sub-Saharan Africa from 2018 through 2030 when targeting geographically according to historical cholera burden and risk factors. We assessed the number of averted cases, deaths, disability-adjusted life-years, and cost-effectiveness with models that account for direct and indirect vaccine effects and population projections over time. Under current vaccine supply projections, an approach optimized to targeting by historical burden is projected to avert 828,971 (95% CI: 803,370-859,980) cases (equivalent to 34.0% of projected cases; 95% CI: 33.2-34.8). An approach that balances logistical feasibility with targeting historical burden is projected to avert 617,424 (95% CI: 599,150-643,891) cases. In contrast, approaches optimized for targeting locations with limited access to water and sanitation are projected to avert 273,939 (95% CI: 270,319-277,002) and 109,817 (95% CI: 103,735-114,110) cases, respectively. We find that the most logistically feasible targeting strategy costs $1,843 (95% CI: 1,328-14,312) per DALY averted during this period and that effective geographic targeting of OCV campaigns can have a greater impact on cost-effectiveness than improvements to vaccine efficacy and moderate increases in coverage. Although our modeling approach did not project annual changes in baseline cholera risk or incorporate immunity from natural cholera infection, our estimates of the relative performance of different vaccination strategies should be robust to these factors.

**Conclusions:** Our study suggests that geographic targeting is critical to the cost-effectiveness and impact of oral cholera vaccination campaigns. Districts with the poorest access to improved water and sanitation are not the same as districts with the greatest historical cholera incidence. While OCV campaigns can improve cholera control in the near-term, without rapid progress in developing water and sanitation services, our results suggest that vaccine use alone are unlikely to allow us to achieve the 2030 goals.

## Introduction

Cholera remains a significant global public health threat, causing more than 100,000 deaths per year globally, with sub-Saharan Africa bearing the majority of the burden [1–3]. In May 2018, the 71st World Health Assembly adopted a resolution aimed at reducing global cholera deaths by 90% by 2030 [4]. Achieving major reductions in morbidity and mortality in sub-Saharan Africa are essential to reaching this goal.

Due to successful campaigns conducted in over 15 countries since 2013 [5], countries that regularly experience cholera are beginning to integrate vaccination with oral cholera vaccine (OCV) into regular public health activities, as recommended by the Global Task Force on Cholera Control (GTFCC) *Roadmap to 2030* and general WHO guidance [6]. These vaccines have 49-67% efficacy in protecting vaccine recipients against cholera infection for up to five years [7], thus presenting an important near-term solution to rapidly reducing cholera risk while long-term improvements to safely managed and sustainable water and sanitation services are made. Moreover, killed whole-cell cholera vaccines are known to be very safe; trials for vaccines currently included in WHO stockpiles have not found evidence for vaccine-related adverse events in non-pregnant or pregnant populations [8,9].

Nevertheless, OCV presents several challenges to traditional approaches to vaccine deployment. OCV does not provide lifelong immunity; while the length of protection is uncertain, it is thought to wane significantly after five years [7]. Further, the vaccine appears to be half as protective in children under five years old [7]. Together, these factors suggest that inclusion of OCV in a childhood vaccination schedule would have limited impact. Similarly, the high degree of clustering of cholera risk, both geographically and demographically [10,11], makes large-scale (e.g., country-wide) vaccination campaigns inefficient.

The limited supply of OCV further complicates its integration into routine cholera control activities. In 2018, 23 million doses of OCV were produced globally (enough to provide the recommended two-dose course to 11.5 million people) [10]. This is only a fraction of what would be needed to cover the 87.2 million people living in high-risk areas of sub-Saharan Africa alone [10]. Consequently, efficient strategies are needed if the limited vaccine supply is to play a significant role in cholera control.

While previous work has examined the impact of targeting OCV to specific age groups [12], an alternate strategy for efficient OCV use is to target vaccines geographically to high-risk areas. Geographic targeting may be particularly important in sub-Saharan Africa, where there is great heterogeneity in cholera dynamics; cholera has an endemic presence in countries like the Democratic Republic of the Congo, Mozambique, and Nigeria, yet causes only sporadic epidemics, separated by multiple years of inactivity, in other places. While cholera outbreaks are often catalyzed by conflict and climate-related events, their magnitude and duration are sustained by limited access to improved water and sanitation, key cholera risk factors.

In this work, we use simulation studies to explore the impact of different approaches to conducting geographically targeted OCV campaigns from 2018 to 2030 in sub-Saharan Africa. We quantify the impact of targeted strategies over untargeted OCV use and compare targeting according to historical cholera burden to that of cholera risk factors. The ultimate aim of these analyses is to provide guidance in how this critical cholera control tool may be used efficiently to accomplish global cholera control goals.

## Methods

### Epidemiologic and Demographic Data Sources

Our methods for mapping cholera incidence have been previously described [10]. Briefly, our estimates of cholera incidence were based on suspected and clinically confirmed cholera case reports from 2010 to 2016 obtained from multiple sources, including the World Health Organization (WHO), Médecins Sans Frontières, ProMED, situation reports from ReliefWeb and other websites, several Ministries of Health, and the scientific literature [3,10]. These cholera reports were combined with ecological risk factors such as access to improved drinking water and sanitation and distance to the nearest major body of water to estimate average annual cholera incidence rates at the 20 km × 20 km grid resolution in a Bayesian modeling framework [3,10]. We did not obtain water and sanitation data for Botswana, Djibouti, and Eritrea, and these countries were excluded from our analyses. Summaries of all cholera data sets, including 20 km × 20 km resolution estimates of cholera cases and incidence and instructions for requesting access are available at http://www.iddynamics.jhsph.edu/projects/cholera-dynamics.

Population estimates for each administrative area were derived from WorldPop’s population density rasters for Africa [13,14]. The population for each 1 km × 1 km grid cell was fit independently to population estimates and projections from WorldPop for 2000, 2005, 2010, 2015, and 2020 using a log-linear model. Annual population estimates from 2018 through 2030 were then projected for each cell independently using the fit log-linear rate of population change.

### Model Simulation

We chose to use a phenomenological model to deal with incidence and the impact of immunity on transmission, as classical mechanistic transmission models (e.g., SIR models) tend to perform poorly on coarse time scales and overestimate case counts while masking underlying heterogeneity in populations.

Our models were simulated in a deterministic manner at a 5 km × 5 km grid resolution. We chose this grid resolution as a balance between computational feasibility and accurate district-level population sizes (as summed across corresponding grid cells); 20 km × 20 km grid resolutions provided highly inaccurate district population estimates, which consequently affected our vaccine targeting exercise. The input cholera incidence rate estimates were disaggregated from a 20 km × 20 km to 5 km × 5 km grid (5 km × 5 km rates were the same as the rate for the corresponding 20 km × 20 km cell), and the WorldPop projections for a given year were aggregated from a 1 km × 1 km to 5 km × 5 km grid.

The uncertainty expressed in the reported confidence intervals reflect the statistical uncertainty captured in the posterior distribution of mean annual incidence across sub-Saharan Africa [10]. We used the same fixed set of 1000 posterior draws of mean annual incidence for all years, vaccine deployment strategies, and sensitivity analyses.

### Vaccine Properties

#### Campaign Coverage

We conducted a review of published literature on post-OCV campaign vaccination coverage surveys and identified seven studies related to 24 two-dose campaigns conducted globally from 2003 through 2016 (Table S1) [15–23]. For each of the seven studies, we resampled two-dose (the standard vaccine regimen) coverage estimates 5000 times from a Gaussian distribution with a mean equal to the estimated coverage at the campaign site and the variance derived from the associated 95% confidence intervals; for studies with multiple locations, we first drew a single location randomly and then sampled from a Gaussian distribution of coverage estimates for that location. We pooled these 35,000 draws across studies and used the median (68%) of the samples as the baseline coverage estimate for our model (Figure S1). These estimates give studies equal weight regardless of the number of campaign locations and campaign site sample size.

#### Vaccine Efficacy

We fit a log-linear decay function to two-dose vaccine efficacy data reported zero to five years after vaccination in a recent meta-analysis [7], and used the mean point estimates for each year as (direct) vaccine efficacy in our model. In this framework, the initial vaccine efficacy is 66% declining to 0% after six years (Figure S2). We modeled vaccination with only the full two-dose regimen with no wastage.

#### Vaccine Indirect Effects

OCV has been shown to induce indirect protection across multiple settings [23–25]. We modeled indirect protection as a function of the vaccination coverage in a given grid cell using data from trials in India and Bangladesh [24,25]. Specifically, the phenomenological association between the relative reduction in incidence among unvaccinated (placebo) individuals and OCV coverage in their ‘neighborhood’ was fit to a logistic function (Figure S3). Under this model of indirect vaccine protection, individuals not protected by vaccine and residing in grid cells with 50% and 70% vaccination coverage experienced an 80% and near 100% reduction in cholera risk compared to no vaccination scenarios, respectively.

### Vaccine Supply Projections

Current global supplies of OCV are limited. In 2017, approximately 17 million bivalent killed whole-cell OCV doses of ShanChol (Shantha Biotech, Hyderabad, India) and Euvichol-Plus/Euvichol (Eubiologics, Seoul, Republic of Korea) were produced. Based on estimates from experts within the GTFCC and data from vaccine manufacturers in the first half of 2018, we assumed that global OCV supply would increase linearly from 23 million doses in 2018 to 59 million doses in 2030 (Figure S5).

### Vaccination Deployment Strategies

We modeled eight OCV deployment strategies: untargeted distribution, four historical burden-based strategies (rate optimized, rate-logistics optimized, case optimized, and case-logistics optimized) (Figure 1), and three based on access to improved water and sanitation (water optimized, sanitation optimized, and watsan optimized). For all targeted strategies, a given district was targeted fully (i.e., achieving 68% vaccination coverage) only once every three years, as suggested by WHO guidelines for OCV deployment [26]. Districts were ranked only once and vaccination campaign targets changed annually according to parameters on vaccination campaign frequency and vaccine supply.

**Figure 1.**
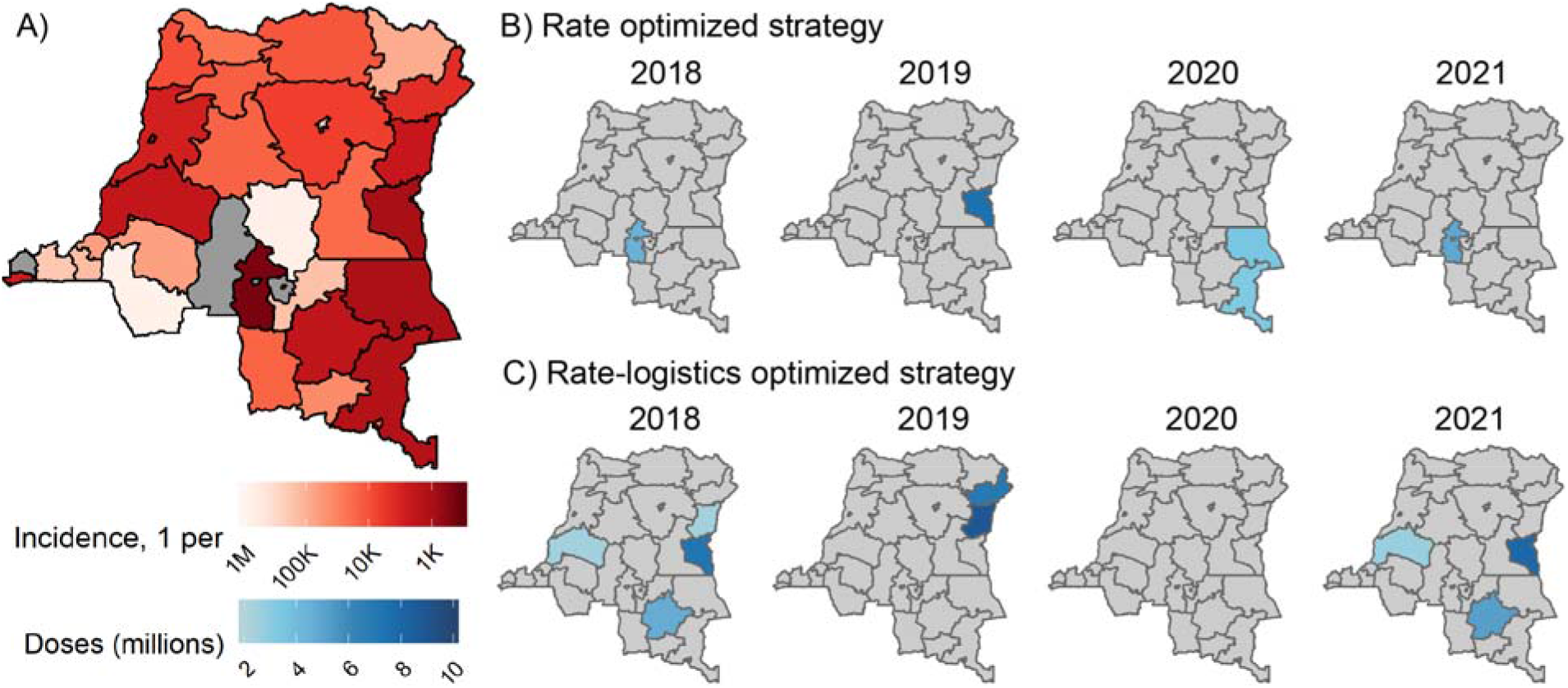
Demonstration of district-level vaccination deployment strategies in the Democratic Republic of the Congo (DRC). A) Estimated annual incidence rate by district from 2010-2016. Districts (International Organization for Standardization second-level, sub-national administrative unit) in grey had an annual incidence rate less than one per million people. Vaccine allocations in the DRC by year according to the B) rate optimized and C) rate-logistics optimized strategies. Districts were targeted in a second consecutive year if they first year’s campaign did not have enough vaccine to cover the target population. Districts in grey were not targeted in DRC that year and there were no districts targeted in the DRC in 2020 in the rate-logistics optimized strategy.

#### Untargeted distribution

OCV is distributed proportional to population throughout the study region (i.e., everyone has equal likelihood of receiving vaccine).

#### Rate optimized

We ranked all districts across countries in the study area by estimated cholera incidence rate (i.e., cases per unit population) and targeted them in decreasing order with vaccine until the annual global vaccine supply was depleted (Figure S7).

#### Rate-logistics optimized

It may not be logistically feasible to target districts in multiple, potentially geographically non-adjacent countries at once. Thus, in the rate-logistics optimized strategy, we first ranked countries according to the size of the population residing in high-risk districts, and then targeted vaccines to all high-risk districts in those countries. The highest risk districts were defined as those where cholera incidence exceeded a threshold of 1 case per 1000 persons for at least 100,000 residents or 10% of the population; subsequent tiers of “high-risk districts” employed incidence thresholds of 1 case per 5000, 10,000, and 100,000 persons successively (Figure S8) [3]. Within each incidence threshold tier, districts were ranked from highest to lowest by estimated cholera incidence rate (hence, rate-logistics). Some districts with high incidence rates may not meet the definition of “high-risk district” because fewer than 100,000 people or less than 10% of the district population reside in cells achieving the given incidence threshold tier; this can lead to differences in targeting between the rate optimized and rate-logistics optimized strategies.

#### Case optimized and Case-logistics optimized

These are the same as their rate optimized counterparts, except districts were ranked according to raw cholera case numbers instead of estimated cholera incidence rates (Supplement only).

#### *Water optimized*, *Sanitation optimized*, and *Watsan optimized*

We ordered districts across all countries according to their estimated coverage of access to improved water, improved sanitation, and improved water or sanitation (“Watsan”, Supplement only), and then targeted them from worst to best coverage. District-level coverage was derived from previously modeled estimates of access to improved water and improved sanitation [27].

### Measuring Public Health Impact and Costs

We estimated the public health impact of conducting OCV campaigns from 2018 to 2030 as the number of cholera cases, deaths, and disability-adjusted life-years (DALYs) averted over the period from 2018 to 2030 (See Supplementary Material for details).

We reviewed four cost surveys for mass OCV campaigns that reported the vaccine delivery and vaccine procurement costs per fully vaccinated person (i.e., receiving two vaccine doses) in non-refugee African settings [28–31] (Table S2). One outlying data point with high vaccine purchase prices ($10 in 2009 USD) was excluded [31], as ongoing policy discussions suggest that future, preventive use of OCV in these settings will be likely available at lower prices. All costs were adjusted to 2017 US dollars (USD) according to the World Bank Consumer Price Index. The mean total cost per fully vaccinated person ($6.32), calculated as the sum of delivery and procurement costs was used to calculate vaccination campaign costs.

Program costs were measured as the cost per DALY averted (2017 USD) and discount rates for health benefits and costs were set to 0% and 3%, respectively [32]. We calculate total costs per DALY averted for all combinations of the three cost estimates and 1000 samples of averted DALYs (per model) in order to estimate 95% confidence intervals jointly from these two empirical distributions.

### Sensitivity Analyses for Model Parameters

We performed one-way sensitivity analyses of differences in vaccination deployment strategy, vaccine efficacy, indirect vaccine protection, vaccination campaign coverage, vaccine supply, net loss of vaccinated individuals due to migrations and deaths (i.e., population turnover rate), and the time between vaccination campaigns in a given location (Table 1). Sensitivity parameters for direct vaccine efficacy were taken directly from the upper and lower 95% confidence interval bounds from a recent meta-analysis (Figure S2) [7]. For indirect vaccine protection, we assumed, as a lower bound, that the vaccine conferred no indirect protection (Figure S4). On the upper end, we assumed that individuals not directly protected by vaccine residing in grid cells with 30%, 50%, and 70% vaccination coverage experienced a respective 66%, 88%, and 97% reduction in cholera risk relative to a no vaccination scenario, according to a logistic model fit to published estimates (Figure S4) (Longini et al. 2007). Sensitivity parameters for vaccination coverage were taken from the 10th and 90th percentile resampled distribution of published coverage survey estimates from previous OCV campaigns (Figure S1, Table S1) [15–23]. Sensitivity parameters for vaccine supply had, after 2019, either no growth or linear growth to 95 million doses in 2030; upper limit vaccine supplies may be achieved if new OCV production facilities open as planned (Figure S6). We explored the sensitivity of our primary estimates to assumptions of the population turnover rate using data from the 2017 UN World Population Prospects Projections. The upper and lower bounds of population turnover rate were represented by the 5th and 95th percentiles of life expectancy for African countries in our study from 2018-2030, 56 and 70 years old, respectively [33].

**Table 1.** Parameters and references for the primary model vaccination campaign performance and costs. Sensitivity analysis parameters are also reported for vaccine efficacy, indirect vaccine protection, vaccination coverage, vaccine supply, population turnover rate, and periodicity of vaccination campaigns.

| **Model parameter** | **Primary Scenario** | **Sensitivity Lower Bound** | **Sensitivity Upper Bound** |
| --- | --- | --- | --- |
| Vaccine efficacy (% after 1-5 years) [7] | 60, 52, 43, 32, 20 | 49, 44, 29, 4, 0 | 68, 59, 54, 52, 51 |
| Indirect vaccine protection (% at 10, 30, 50, 70, 90% coverage) [24,25,42] | .09, 7, 84, 100, 100 | 0, 0, 0, 0, 0 | 34, 66, 88, 97, 99 |
| Vaccination coverage (%) [Refs in Table S1] | 68 | 50 | 84 |
| Vaccine supply (2018-2030 in millions) | 23-59 | 23-26 | 23-95 |
| Population turnover rate (years<sup>-1</sup>) [33] | 65 | 70 | 56 |
| Periodicity of vaccination campaigns | Every 3 years | Every 5 years | Every 2 years |
| Vaccine delivery cost per FVP (2017 USD)<sup>1</sup> [Refs in Table S2] | 2.33 | Not applicable | Not applicable |
| Vaccine procurement cost per FVP (2017 USD)<sup>2</sup> [Refs in Table S2] | 5.49 | Not applicable | Not applicable |

<sup>1</sup> Vaccine delivery costs are related to program preparation, administration, and adverse events following immunization.
<sup>2</sup> Vaccine procurement costs are related to vaccine price, shipment, and storage.

### Projected Cases Averted Due to Vaccination Campaigns

For our study period years of 2018-2030, we assumed that cholera incidence remained constant at the mean annual incidence rates observed from 2010-2016 in the absence of vaccination. Exploratory analyses using annual cholera reports to the WHO suggest that mean annual incidence may be an unbiased estimator for annual incidence at the country-level, which suggests that this assumption about projected cholera incidence is valid in the expectation (Figure S11), although uncertainty remains under-estimated.

We assume that cholera vaccination is the only mechanism that confers immunity to cholera. The proportion of the population not protected by vaccine, and hence susceptible to cholera infection (‘susceptibles’), in location *i* in year *t* is:

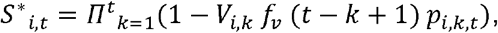

where *V*_*i,k*_ is the proportion of the population vaccinated in year *k* at location *i*. The function *f*_*v*_ (*n*) represents the direct vaccine efficacy *n* years after vaccines were administered to the population. We modeled demographic changes in the population as

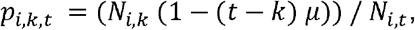

where *p*_*i,k,t*_ is the proportion of the population in location *i* from year *k* that is still present in year *t*. This proportion is calculated as a function of the population *N* in years *k* and location *i*, and the location’s net loss of vaccinated individuals due to migrations and deaths *μ*. We assumed that the net loss of vaccinated individuals due to migrations and deaths (population turnover rate) was the same for all locations and tied roughly to median life expectancy across African countries in our data from 2018-2030 (65 years^−1^) [33]. The expected number of cholera cases at time *t* and location *i*, *Y*_*i,t*_, is then calculated as

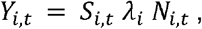

where *S*_*i,t*_ is the susceptible proportion of the population after accounting for both direct and indirect vaccine effects, and *λ*_*i*_ is the projected baseline cholera incidence for location *i*. The total proportion of effective susceptibles *S*_*i,t*_ may be represented as *S*_*i,t*_ = *S*\*_*i,t*_ × *g*_*v*_ (*S*\*_*i,t*_), where *g*_*v*_ is a function that models the indirect effects of vaccination, as described in the ‘Vaccine Properties’ section (Figures S3). To estimate the potential reductions in cases attributable to OCV use, we calculated

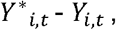

where *Y*\*_*i,t*_ represents the counterfactual scenario where no vaccines were deployed.

#### Data and Code Availability

Model code, input data, processed model outputs have been made available on Github at: https://github.com/HopkinsIDD/geotargeting_ocv_ssa. Raw model output files are available upon request.

## Results

Absent substantial changes to cholera prevention and control, or secular trends in incidence, we expect 2.44 (95% CI: 2.41-2.48) million reported cholera cases from 2018 through 2030 in sub-Saharan Africa; with nearly 50% of these cases in just four countries -- the Democratic Republic of the Congo, Nigeria, Somalia, and Sierra Leone. Given this clustering of disease burden, we examined practical deployment strategies targeting the highest risk districts across sub-Saharan Africa according to historical cholera burden (rate optimized and rate-logistics optimized) and to water and sanitation coverage (water optimized and sanitation optimized) and assessed the sensitivity of our results to different vaccine-related assumptions.

When targeting districts ranked by expected cholera incidence rate (rate optimized), 34.0% (95% CI: 33.2-34.8%) of cases that would have otherwise occurred without vaccination from 2018 through 2030 were averted (Figure 2). This reduction translates to 828,971 (95% CI: 803,370-859,980) cases, 31,958 (95% CI: 31,503-33,011) deaths, and 746,749 (95% CI: 736,607-762,273) DALYs averted after vaccination campaigns from 2018 through 2030 (Figures S13-S15, Table S4). Due to our model assumption that baseline cholera risk will remain constant over the study period, the rate optimized strategy represents the “best-case” district targeting scenario, and results from targeting strategies should be interpreted relative to one another.

**Figure 2.**
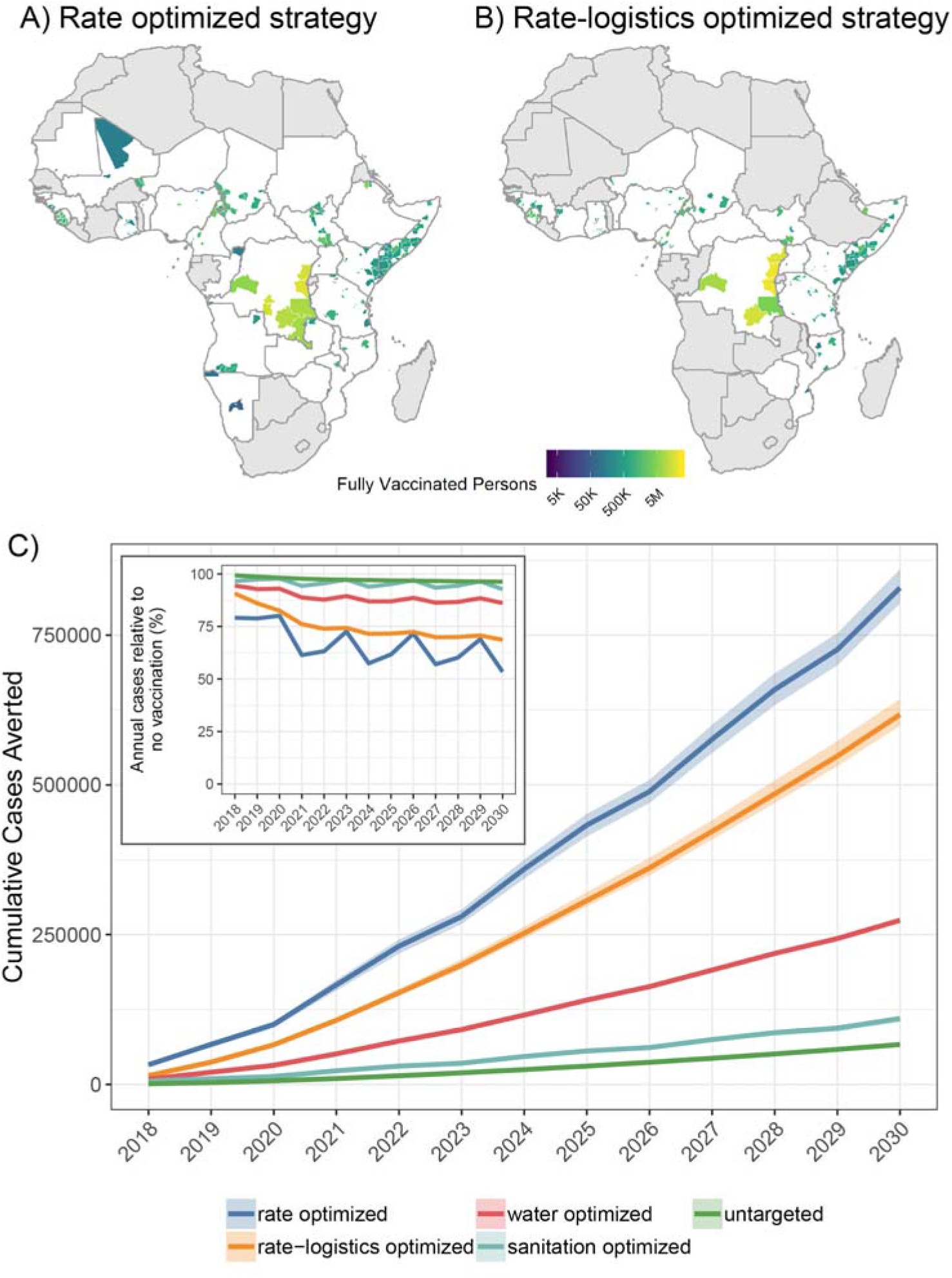
Health outcomes after vaccination under primary model assumptions. Cumulative number of fully vaccinated persons in sub-Saharan Africa as a result of campaigns from 2018 through 2030 according to the A) rate optimized and B) rate-logistics optimized vaccination deployment strategies. Countries in grey had no districts targeted by a given vaccination deployment strategy. C) Cumulative cases averted from mass oral cholera vaccination campaigns across five deployment strategies in sub-Saharan Africa from 2018 through 2030 (mean and 95% CI). The inset figure shows the mean annual percentage of cholera cases averted in our models according to each deployment strategy.

Taking a logistically simpler approach to incidence rate based targeting, where high-risk districts within high-risk countries are targeted together (rate-logistics optimized), 25.3% (95% CI: 24.8-26.1%) of cases that would have otherwise occurred without vaccination were averted cumulatively, translating to 617,424 (95% CI: 599,150-643,791) cases, 24,189 (95% CI: 23,579-25,115) deaths, and 577,533 (95% CI: 564,572-590,935) DALYs averted after 13 years of vaccination campaigns (Figure 2, Figures S13-S15, Table S4).

Targeting districts geographically by lack of access to improved water and sanitation was not as effective as targeting by historical cholera disease burden. When targeting districts by lack of access to improved water (water optimized), 11.2% (95% CI: 11.1-11.4%) of cases were averted; this translates to 273,939 (95% CI: 270,319-277,002) cases, 10,672 (95% CI: 10,517-10,827) deaths, and 255,090 (95% CI: 251,723-258,787) DALYs averted after 13 years of vaccination campaigns (Figure 2, Figures S13-S15, Table S4). Targeting by lack of access to improved sanitation was substantially less effective (sanitation optimized); 4.5% (95% CI: 4.3-4.6%) of cases were averted, representing 109,817 (95% CI: 103,735-114,110) cases, 3,682 (95% CI: 3,469-3,812) deaths, and 83,228 (95% CI: 78,579-86,117) DALYs that would have otherwise occurred without vaccination from 2018 through 2030 (Figure 2, Figures S13-S15, Table S4).

Across all years with vaccination campaigns, the rate optimized strategy averted 20-47% of annual cholera cases (range of mean estimates across years from 2018 to 2030) and the rate-logistics optimized strategies averted 9-31% of annual cholera cases (Table S5). Burden based deployment strategies substantially outperformed the water optimized and sanitation optimized strategies, which averted 6-14% and 2-7% of annual cholera cases, respectively (Table S5). The *untargeted* strategy, where vaccine was deployed at equal coverage across all districts, saw a 0.7-4% annual case reduction from 2018 to 2030 (Table S5).

The most effective vaccination deployment strategies were also the most cost-effective in our simulations. We projected mean costs of $1,843 (95% CI: 1,032-2,382) and $2,383 (95% CI: 1,327-3,102) per DALY averted (USD 2017) for the rate optimized and rate-logistics optimized strategies, respectively (Figure 3, Table S4). Targeting by risk factors was much more expensive; mean costs for the water optimized and sanitation optimized strategies were $5,394 (95% CI: 3,029-6,965) and $16,546 (95% CI: 9,121-22,243) per DALY averted (USD 2017), respectively (Table S4). As a point of reference, the 2017 gross domestic products (GDPs) of countries within our study area ranged from roughly $300 to $10,000, with a mean around $1,734 (2017 USD); interventions are typically defined as cost-effective if the mean cost per DALY averted is less than 3 times the GDP of a country and highly cost-effective if it is less than or equal to the GDP of a country [32,34].

**Figure 3.**
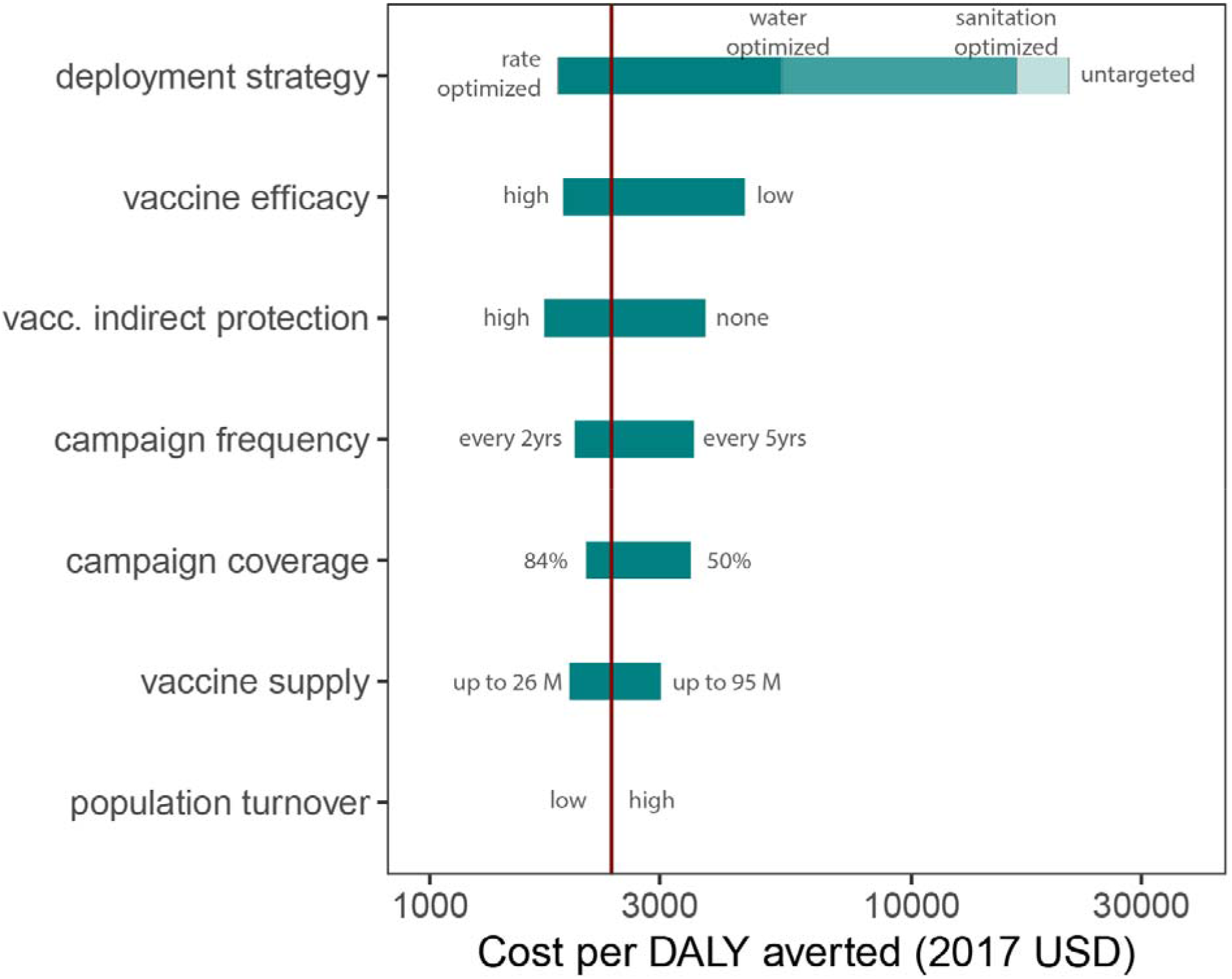
Sensitivity analysis of the mean cost per DALY averted to alternate parameters for vaccination deployment strategy, vaccine efficacy, indirect protection from vaccine, vaccination campaign frequency, vaccination coverage, vaccine supply, and population turnover rate. The red vertical line indicates the mean cost per DALY averted for the *rate-logistics optimized* scenario with the primary model parameters ($2,383). The *untargeted* and *rate optimized* strategies represented the highest and lowest cost vaccination deployment strategies, respectively.

We examined the one-way sensitivity of our results to alternate parameters for vaccine efficacy, indirect protection from vaccine, vaccination campaign frequency, vaccination coverage, vaccine supply, and population turnover, and campaign deployment strategies when taking the rate-logistics optimized strategy as the primary scenario (Table 1, Figure 3). Changing the vaccination deployment strategy affected the greatest variation in mean cost per DALY averted, from $1,843 (95% CI: 1,032-2,382) to $21,213 (95% CI: 11,953-27,263) for the rate optimized and untargeted strategies, respectively (Table S4). Vaccine efficacy affected the second-largest variation among model parameters, where the 97.5 and 2.5 percentile vaccine efficacy estimates (Table 1) from a recent meta-analysis had mean costs per DALY averted of $1,892 (95% CI: 1,053-2,463) and $4,507 (95% CI: 2,512-5,864), respectively (Tables S6-S7). The model was nearly as sensitive to assumptions about indirect vaccine protection, where assumptions of high indirect protection and no indirect protection (Table 1) had mean costs per DALY averted of $1,729 (95% CI: 963-2,250) and $3,733 (95% CI: 2,079-4,859), respectively (Tables S8-S9). Model results were less sensitive to vaccination campaign frequency, vaccination coverage, vaccine supply, and population turnover rates; these results are reported in Tables S10-S17.

## Discussion

This study shows the essential role of geographic targeting in guaranteeing that extended cholera vaccination campaigns across sub-Saharan Africa would have a measurable impact on cholera incidence. When considering the direct and indirect protective effects of oral cholera vaccines, our results suggest that, under projected resource constraints, campaigns that are geographically targeted according to disease burden may avert over 8 times more cholera cases, deaths, and DALYs than untargeted (i.e., general population) approaches. Vaccination deployment strategy can have a greater impact on health impact and cost-effectiveness than substantial improvements to vaccine efficacy, vaccination campaign frequency, and vaccination coverage.

Recent discussions and guidance by the Global Task Force on Cholera Control have suggested that priority areas for preventive cholera control (burden hotspots) should be ranked according to mean annual incidence, and that risk factors like access to water and sanitation should have a secondary role. Decision making around OCV allocation and targeting is complex and different actors are concerned with maximizing public health impact, navigating global and local politics, and considering the ethics of balancing preventive, responsive, and humanitarian uses of vaccine. Our results contribute to the discussion around maximizing the public health impact for preventive and potentially routine OCV campaigns, but they may not be best targeted to locations at risk for new cholera introductions. Seventy percent of the targets in the best-performing strategy are in countries with endemic cholera, and it is likely that reactive campaigns would occur in locations with less predictable cholera patterns (Table S3). While it is unlikely that OCV targeting decisions will occur on a continental scale in the real world, we sought to examine the epidemiological principles that will likely be considered in future OCV policy discussions.

Two vaccination deployment strategies with different disease burden-based criteria for identifying high-risk districts in Africa yielded similarly effective results. We believe the rate-logistics optimized approach to be the most practical deployment strategy considered. In this approach, countries are ranked by population living in high-risk districts and then only these high-risk districts are targeted for vaccination. While the less practical rate optimized strategy (which prioritized districts based on burden regardless of country) does prevent more cases, the resulting benefit would not likely outweigh the unmeasured, added costs in logistical implementation. In targeting the highest risk geographic locations, our study suggests that vaccination can be a cost-effective cholera treatment strategy, thus complementing more comprehensive analyses that suggest the importance of targeting high-risk demographic groups for cost-effective cholera vaccination [35].

Improved access to safe water and sanitation is necessary for long-term cholera control and reductions in overall diarrheal disease burden [36–38]. We examined the impact of vaccination deployment strategies that targeted districts with the lowest access to improved water and sanitation as measured by WHO/UNICEF Joint Monitoring Programme for Water Supply, Sanitation and Hygiene (JMP) indicators [27,39]. While our model assumptions guaranteed the superior performance of the rate optimized strategy, districts with the poorest access to improved water and sanitation were not the same as those with the highest historical cholera burden. Targeting by water and sanitation access for the same number of deployed vaccines averted less than one-third as many cholera cases as targeting by disease burden-based criteria (Table S4). However, our analyses were based on indicators that have been criticized for their focus on access to water and sanitation infrastructure as opposed to safely managed and sustainable water and sanitation use [40], which may explain the relatively poor performance of the water and sanitation optimized strategies. The JMP recently adopted new measurement criteria, which may prove to be more specific indicators of cholera risk [41].

We examined the sensitivity of our model to different assumptions about cholera and cholera vaccine dynamics in our models, but there remain several limitations outside the scope of these analyses (Tables S6-S17). Our models do not account for immunity due to natural cholera infection, and the indirect effects of vaccination are captured only at the grid-cell level, which limits the estimated impact of the vaccination at critical hubs of cholera transmission. Our projections assume that baseline cholera incidence (measured as mean annual incidence) remains constant throughout the study period, which may not capture secular trends, or inter-year variability in cholera incidence or surveillance, or the emergence of new cholera patterns due to conflict, crisis, or changing epidemiology. However, mean annual incidence appears to be an unbiased estimator for annual incidence (Figure S11), suggesting that our results are valid in the expectation although uncertainty remains under-estimated. Additionally, our baseline incidence estimates represent only reports of suspected cholera cases, making no explicit adjustments for biased reporting or measurement error. Further, we examined only the impact of two-dose vaccination campaigns; recent evidence suggests that a single OCV dose may be efficacious [7,40–42], but no data exist on its long-term protection relative to a two-dose course. While the inclusion of age-specific parameters in our model might be biologically justified, there is a dearth of age-specific cholera incidence data and long-term vaccine efficacy estimates in children [7].

Improvements to the coverage and geographic resolution of cholera surveillance and short-term migration patterns, and an improved understanding of the relationship between cholera, climate, and disruptive events may enable future versions of this model to characterize optimal OCV targeting for epidemic and endemic settings. Future water, sanitation, and hygiene (WASH) survey data collected under the new JMP definitions, combined with higher quality cholera surveillance data, may make nuanced treatment of complex epidemiological interactions possible. For instance, projections of OCV impact could employ more complex targeting strategies that account for interactions between cholera immunity due to recent outbreaks and probability of epidemic cholera introduction and propagation due to limited WASH access.

Our results show how geographic targeting can play an essential role in ensuring that mass OCV use leads to substantial reductions in the global burden of cholera, even under current supply constraints. Strategic targeting of resources can play an essential role in making cholera control efforts cost-effective, a message that may be generalized to the entire suite of cholera control activities, including those to increase access to safe water and improved sanitation. Continued increases in global OCV production would enable a greater proportion of high-risk populations to be targeted with vaccination with greater regularity, but our results suggest that even the most effective geographic targeting strategies paired with optimistic OCV supply projections will not be enough to achieve the global cholera burden reduction goals set forth by the World Health Assembly resolution and the *Roadmap to 2030*. Substantial improvements to other sectors of the Roadmap, including broad investments and progress to improve water and sanitation infrastructure and cholera surveillance, will be needed to make headway in this ambitious initiative.

## Contributors

ECL, ASA, and JL designed and conceived the study, interpreted data, and wrote the manuscript. ECL analysed data and prepared the figures. JK contributed code and facilitated data management. JK, SMM, and HSM supervised data entry and integrity. SMM developed baseline maps of cholera incidence. All authors provided writing input and reviewed the final draft.

## Acknowledgments

We would like to acknowledge the Cholera Team at WHO, including Lorenzo Pezzoli and Dominique Legros for their input on critical parameters in our model, Didier Bompangue, Abdinasir Abubakar, Epicentre/MSF, UNICEF, and the Ministries of Health of Benin, Cameroon, the Democratic Republic of the Congo, Malawi, Mozambique, Nigeria, South Sudan, Uganda, Zambia, and Zimbabwe for sharing data, and Melissa Ko, on behalf of Gavi, for her feedback on estimating the health impacts of OCV.

