## Supplementary Material for "The impact of geographic targeting of oral cholera vaccination in sub-Saharan Africa: a modeling study"

*Elizabeth C. Lee, Andrew S. Azman, Joshua Kaminsky, Sean M. Moore, Heather S. McKay, Justin Lessler*

### Vaccination Campaign Coverage

We assessed the sensitivity of our models to vaccination campaign coverage, as there has been great variability in the success of previous OCV campaigns. We first conducted a review of published literature on post-OCV campaign vaccination coverage surveys and identified seven studies related to 24 two-dose campaigns conducted globally from 2003 through 2016 (Table S1) (Lam et al. 2017; Tohme et al. 2015; Luquero et al. 2013; Uddin et al. 2014; Massing et al. 2018; Sema Baltazar et al. 2018; Cavailler et al. 2006).

Table S1: **Published coverage survey estimates from previous OCV campaigns.** Median point estimates and standard errors were used to resample coverage from a normal distribution for each campaign.

| Study | Campaign | Coverage | SE | Sample Size |
|---|---|---|---|---|
| Lam 2017 | Dahuk 2-dose, Iraq | 0.900 | 0.0255102 | 390 |
| Lam 2017 | Erbil 2-dose, Iraq | 0.930 | 0.0153061 | 512 |
| Lam 2017 | Sulaymaniya 2-dose, Iraq | 0.930 | 0.0178571 | 552 |
| Lam 2017 | Anbar 2-dose, Iraq | 0.980 | 0.0153061 | 550 |
| Lam 2017 | Wasit 2-dose, Iraq | 0.910 | 0.0255102 | 575 |
| Lam 2017 | Salah Addin 2-dose, Iraq | 0.810 | 0.0306122 | 652 |
| Lam 2017 | Najaf 2-dose, Iraq | 0.740 | 0.0357143 | 773 |
| Lam 2017 | Baghdad Karkh 2-dose, Iraq | 0.370 | 0.0382653 | 1273 |
| Lam 2017 | Kerbala 2-dose, Iraq | 0.300 | 0.0408163 | 1763 |
| Lam 2017 | Babil 2-dose, Iraq | 0.210 | 0.0306122 | 2405 |
| Tohme 2015 | Petit Anse 2-dose, Haiti | 0.625 | 0.0229592 | 1118 |
| Tohme 2015 | Cerca Carvajal 2-dose, Haiti | 0.768 | 0.0272959 | 809 |
| Luquero 2013 | Boffa 2-dose, Guinea | 0.780 | 0.0459184 | 840 |
| Luquero 2013 | Douprou 2-dose, Guinea | 0.760 | 0.0408163 | 541 |
| Luquero 2013 | Koba 2-dose, Guinea | 0.690 | 0.0459184 | 935 |
| Luquero 2013 | Mankountan 2-dose, Guinea | 0.840 | 0.0357143 | 576 |
| Luquero 2013 | Tamita 2-dose, Guinea | 0.780 | 0.0510204 | 205 |
| Luquero 2013 | Tougnifili 2-dose, Guinea | 0.770 | 0.0561224 | 826 |
| Luquero 2013 | Kaback 2-dose, Guinea | 0.740 | 0.0357143 | 764 |
| Luquero 2013 | Kakossa 2-dose, Guinea | 0.780 | 0.0459184 | 113 |
| Uddin 2014 | Dhaka second dose, Bangladesh | 0.790 | 0.0267857 | 39910 |
| Massing 2018 | Kalemie Strata 1 2-dose, DR Congo | 0.672 | 0.0257653 | 1159 |
| Sema Baltazar 2018 | Nampula, card and oral report, 2-dose, Mozambique | 0.512 | 0.0673469 | 648 |
| Cavailler 2006 | Beira, 2-dose, Mozambique | 0.536 | 0.0153061 | 82381 |

The study site coverage estimates were skewed slightly left due to the presence of outlying low vaccination campaigns among our study pool (with a mean of 71.3% and median of 76.9%).

For each of the seven studies, we resampled two-dose (the standard vaccine regimen) coverage estimates 5000 times from a Gaussian distribution with a mean equal to the estimated coverage at the campaign site and the variance derived from the associated 95% confidence intervals; for studies with multiple locations, we first drew a single location randomly and then sampled from a Gaussian distribution of coverage estimates for that location. We pooled these 35,000 draws across studies and used the median (68%) of the samples as the baseline coverage estimate for our model (Figure S1). The 10th (49%) and 90th (84%) percentiles of the resampled distribution were used for the low and high coverage sensitivity analyses. These estimates give studies equal weight to the distribution regardless of the number of campaign locations.

As there are many possible methods for identifying OCV coverage parameters for our models, we compared the parameters we used in our study (shown above), to others that could have been proposed. As one alternative, we performed the resampling exercise where weights were applied according to campaign site sample size (Figure S1). The median was 66%, and the 10th and 90th percentiles were 44% and 82% respectively. As a second alternative, we performed the equal-weighting sampling exercise again under the assumption that coverage should follow a lognormal distribution (Figure S1). The median was 68%, and the 10th and 90th percentiles were 49% and 84% respectively.

The percentiles of interest were the same up to the 100th decimal place for the unweighted Gaussian (used in the main text) and lognormal resampling approaches, while the weighted Gaussian resampled percentiles were slightly lower than those from the unweighted Gaussian resampling scheme.

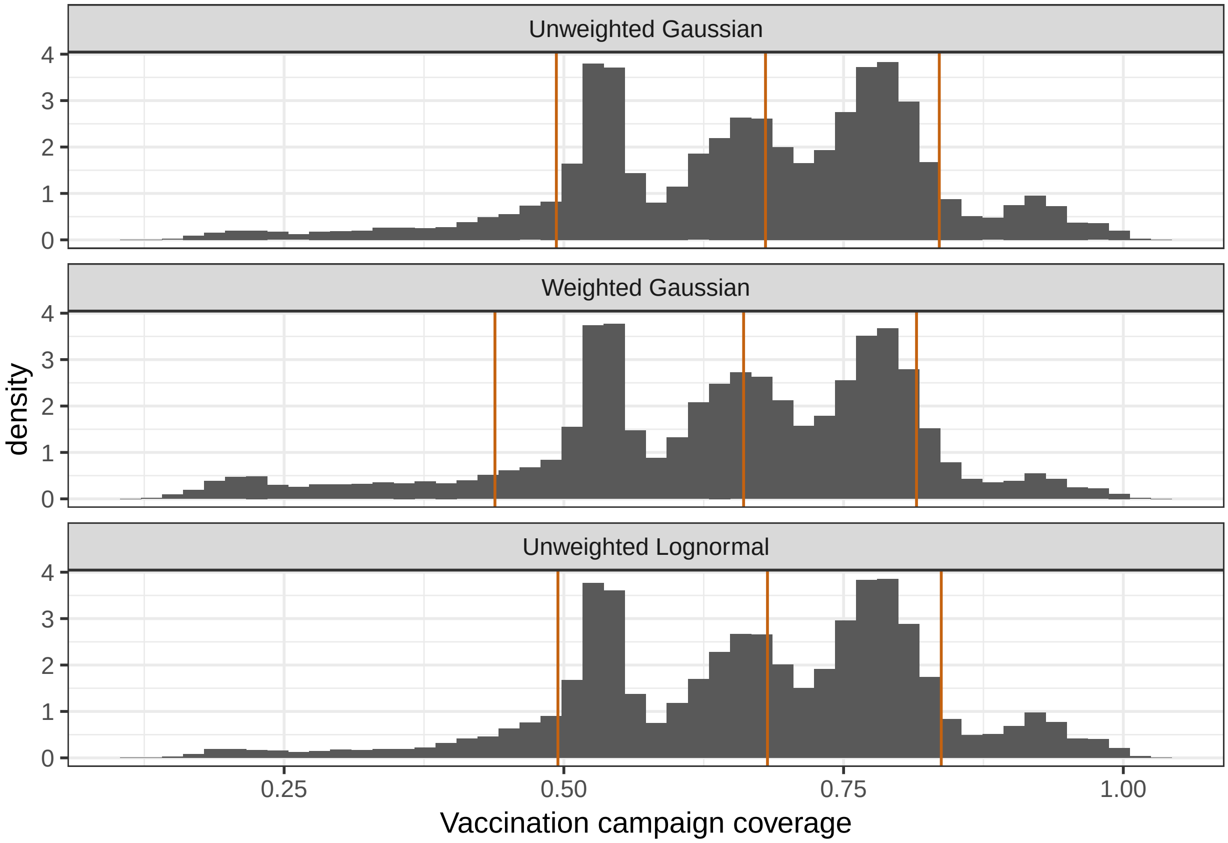

Figure S1: **Resampled distributions of OCV campaign coverage, with Gaussian assumptions and equal weights across campaign sites, which was used for the main paper (top), with Gaussian assumptions and weights for campaign site sample size (middle) and with lognormal assumptions and equal weights across campaign sites (bottom).** We generated a distribution of OCV campaign coverage estimates for each resampling scheme from published OCV coverage surveys. Rust orange lines indicate the 10th percentile, median, and 90th percentile of the resampled distributions.

### Vaccine Efficacy

We examined the sensitivity of our OCV health impact estimates to different vaccine efficacy assumptions. We fit a log-linear decay function to two-dose vaccine efficacy data reported zero to five years after vaccination in a recent meta-analysis (Bi et al. 2017); study estimates were weighted by the inverse of the squared standard error in the model. We then used the mean point estimates for each year as (direct) vaccine efficacy in our model. In this framework, the initial vaccine efficacy is 66% declining to 0% after six years (Figure S2). We modeled vaccination with only the full two-dose regimen with no wastage.

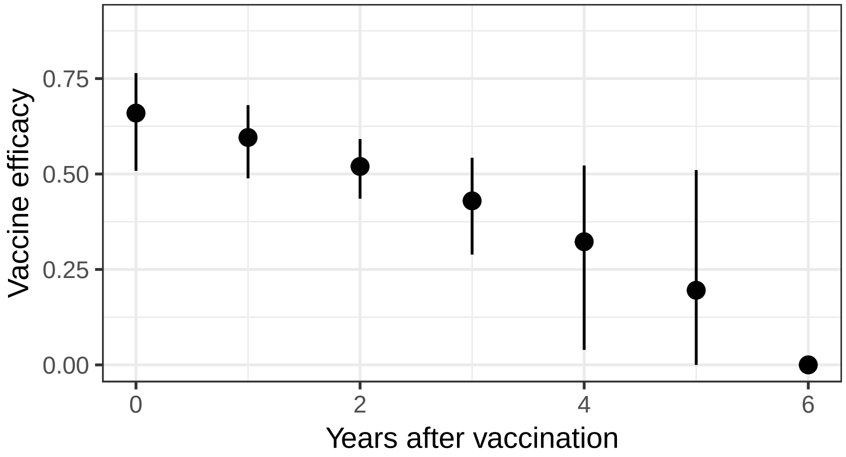

Figure S2: **Waning vaccine efficacy in years after vaccination.** The data displayed represents the mean point estimate and 95% confidence intervals across data collected across seven vaccine efficacy studies. Mean point estimates were used to parametrize the primary model, while the 2.5 and 97.5 percentiles were used to parametrize the low and high vaccine efficacy models, respectively.

### Vaccine Indirect Effects

Due to limited data and uncertainty about the spatial scale of indirect vaccine protection, we assessed the sensitivity of our modeled health impacts to different indirect vaccine protection assumptions. For our baseline parameter set, we modeled indirect protection as a function of the vaccination coverage in a given grid cell using data from trials in India and Bangladesh (Ali et al. 2005, 2013). Specifically, the phenomenological association between the relative reduction in incidence among unvaccinated (placebo) individuals and OCV coverage in their "neighborhood" was fit to a logistic function (Figure S3). Under this model of indirect vaccine protection, individuals not protected by vaccine and residing in grid cells with 50% and 70% vaccination coverage experienced an 80% and near 100% reduction in cholera risk compared to no vaccination scenarios, respectively.

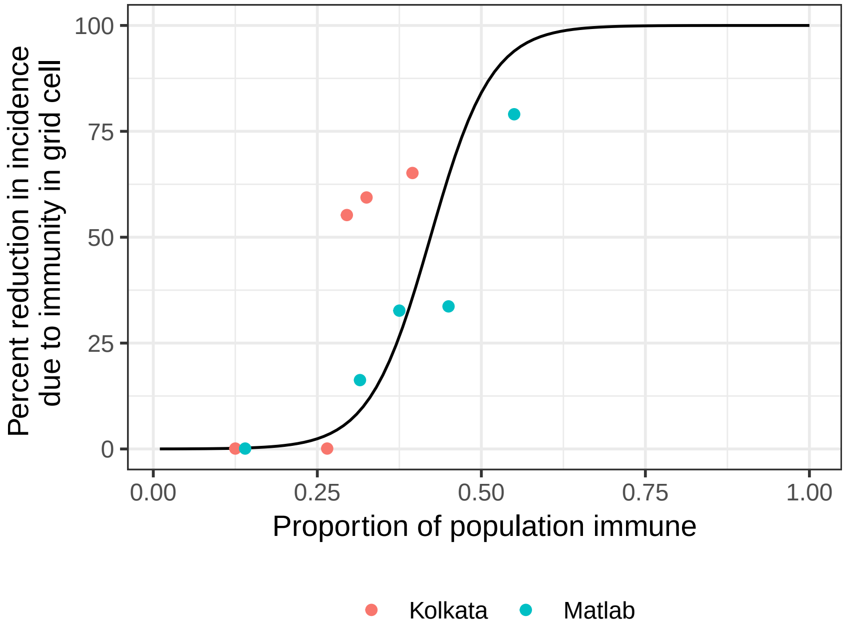

Figure S3: **Indirect protective effect of vaccination.** Each point represents the percent reduction in incidence due to cholera vaccination coverage (population immunity), as reported by randomized controlled trials in Kolkata and Matlab. The black line represents the logistic function fit to these data.

Sensitivity parameters for indirect vaccine effects assumed, at the lower end, that vaccination conferred no indirect protection. On the upper end, we assumed that individuals not protected by vaccine and residing in grid cells with 30%, 50%, and 70% vaccination coverage experienced a respective 66%, 88%, and 97% reduction in cholera risk, according to a logistic model fit of published estimates (Figure S4) (Longini Jr. et al. 2007).

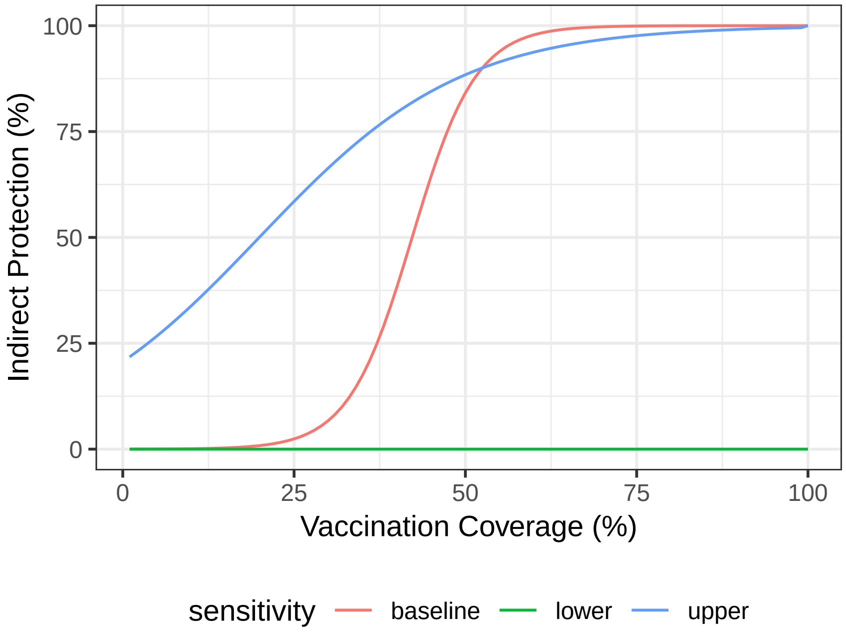

Figure S4: **Sensitivity parameters for indirect protective effect of vaccination.**

### Vaccine Supply Projections

There is great uncertainty in OCV supply over the next decade; increasing demand has led to plans to open several new manufacturing facilities, but the timing of these projected increases in supply remains uncertain. Based on estimates from experts within the GTFCC and data from vaccine manufacturers in the first half of 2018, we assumed that global OCV supply would increase linearly from 23 million doses in 2018 to 59 million doses in 2030 (Figure S5).

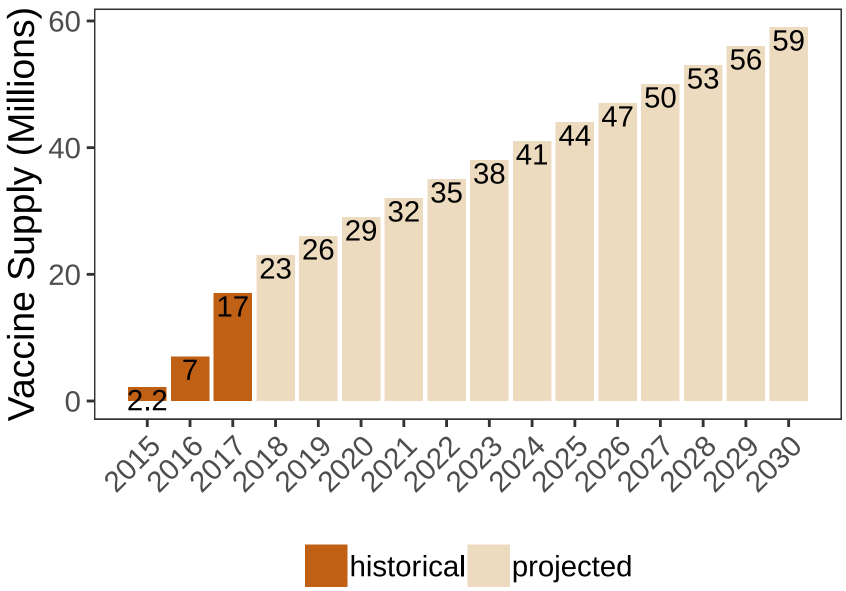

Figure S5: **The historical (2015-2017) and projected (2018-2030) cholera vaccine supply.** The projected vaccine supply per year (in millions) was used as a model input.

Sensitivity parameters for vaccine supply assumed after 2019 either no growth or linear growth to 95 million doses in 2030; upper limit vaccine supplies may be achieved if new OCV production facilities open as planned (Figure S6).

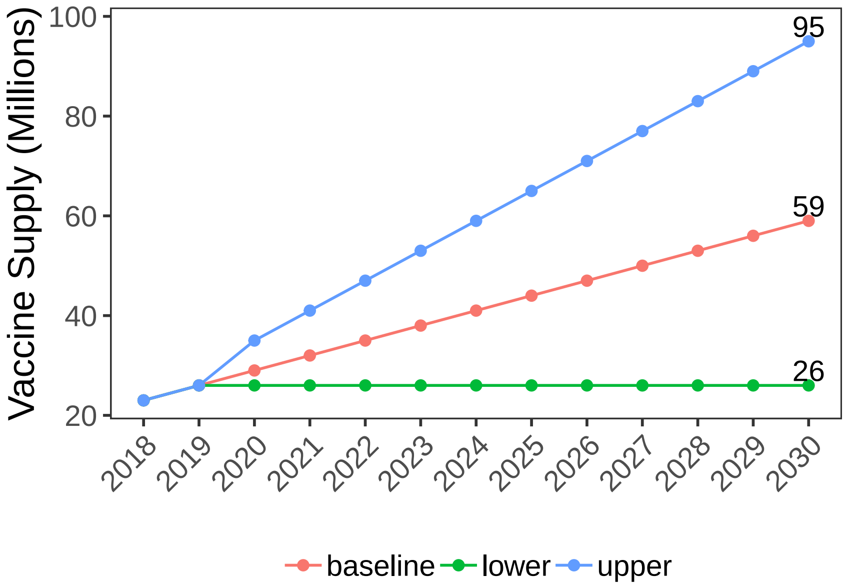

Figure S6: **Sensitivity parameters for projected cholera vaccine supply (in millions) from 2018-2030.**

### Vaccination Deployment Strategies

The case optimized, case-logistics optimized, and watsan optimized results were not reported in the main text. The case optimized strategy performed worse than the rate optimized but better than the rate-logistics optimized strategies; 28.7% (95% CI: 26.6-29.7%) of cases that would have otherwise occurred without vaccination from 2018 through 2030 were averted. This reduction translates to 698,000 cases, 28,000 deaths, and 657,000 DALYs averted after 13 years of vaccination. In the case-logistics optimized strategy, 25.4% (95% CI: 24.5-26.4%) of cumulative cases that would have otherwise occurred without vaccination were averted, thus translating to 619,000 cases, 24,000 deaths, and 581,000 DALYs averted after 13 years of vaccination.

The watsan optimized strategy was similar to the water optimized and sanitation optimized strategies, but the districts were prioritized according to those with the lowest access to improved water or sanitation. As a combination of the water and sanitation optimized strategies, the watsan optimized strategy performed better than the sanitation optimized strategy and less well than the water optimized strategy; 8.2% (95% CI: 7.7-8.4%) of cases that would have otherwise occurred without vaccination from 2018 through 2030 were averted, which translates to 199,000 cases, 7,000 deaths, and 171,000 DALYs averted after 13 years of vaccination campaigns.

The rate optimized and rate-logistics optimized vaccination campaign deployment strategies are described in greater detail in Figure S7 and Figure S8.

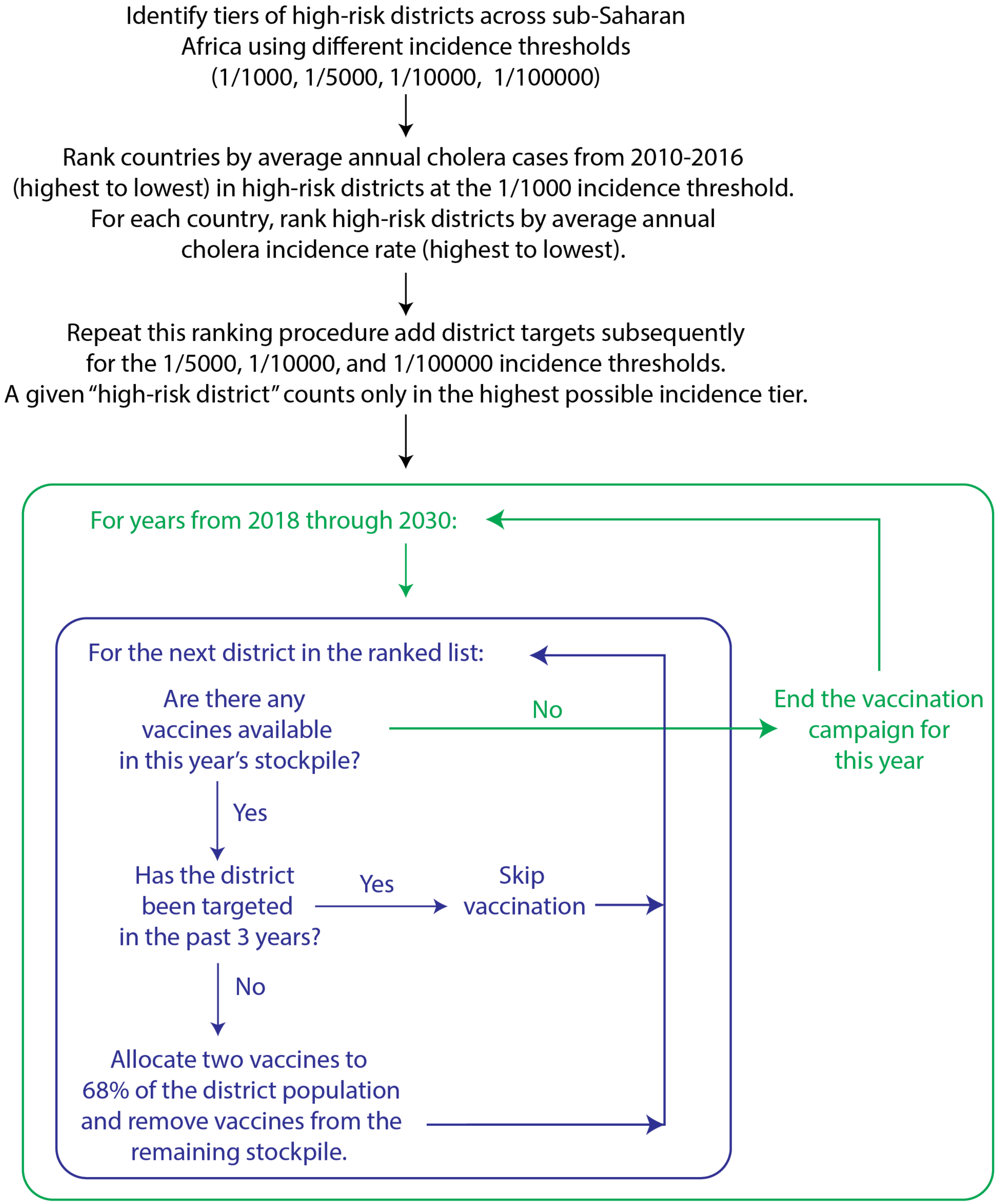

Figure S7: **Flowchart depicting the rate optimized vaccination deployment strategy with primary scenario parameters.**

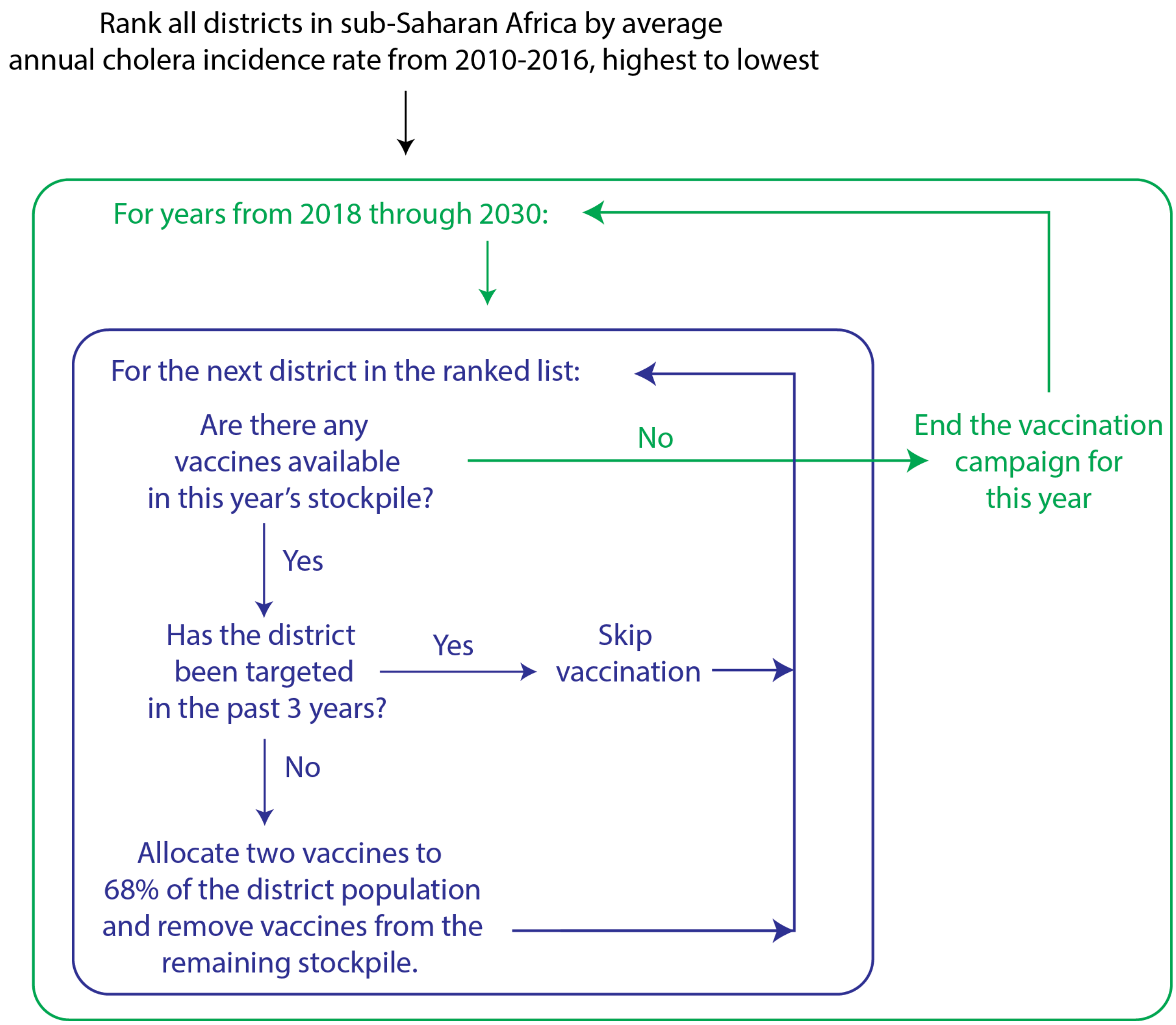

Figure S8: **Flowchart depicting the rate-logistics optimized vaccination deployment strategy with primary scenario parameters.**

### Measuring Public Health Impact and Cost Benefit

We calculated the number of cases averted as the difference in cases between vaccination and no vaccination scenarios. To estimate the number of deaths averted, we multiplied a country-specific case-fatality ratio (CFR) with the expected cases outputs. We estimated country-level CFRs as the ratio of the total number of deaths to cases across all years where cholera cases were reported in the WHO Global Health Observatory database from 1970 to 2016 among countries in our study region (Figure S9). For countries where the estimated CFR exceeded 7% (often based on outbreaks with a small total number of cases), we applied the mean CFR across all remaining countries (3.4%).

We also estimated the disability-adjusted life-years (DALYs) for the vaccination and no-vaccination scenarios, where DALYs are defined as the sum of years of life lost (YLLs) and years of life disabled (YLDs). YLLs were calculated as

$$YLL_i = CFR_i(Y_{i,t} - Y*_{i,t})(\kappa_{i,t-\rho_i} - \rho_i),$$

where $\rho_i$ is the average age of cholera infection in location i and $\kappa_{i,t-\rho_i}$ is the average life expectancy for someone of average cholera infection age in year t. Country-level estimates of life expectancy for each projected year in our study period were obtained from the United Nations (United Nations Secretariat, Department of

Economic and Social Affairs, Population Division 2017) and average age of cholera infection was calculated as the inverse-variance-weighted mean age among cases identified in OCV trials in Africa and the Caribbean (25.75 years old) (Azman et al. 2016; Ferreras et al. 2018; Luquero et al. 2014; Khatib et al. 2012). YLDs were calculated as the product of the proportion of the year disabled with illness (assumed at 4/365) and a disability weight for severe diarrheal disease as estimated from the 2016 IHME Global Burden of Disease Study (0.247) (Institute for Health Metrics and Evaluation, Global Burden of Disease Collaborative Network 2017). Disability weights range from 0 to 1 and represent the magnitude of the health loss associated with a specific health status.

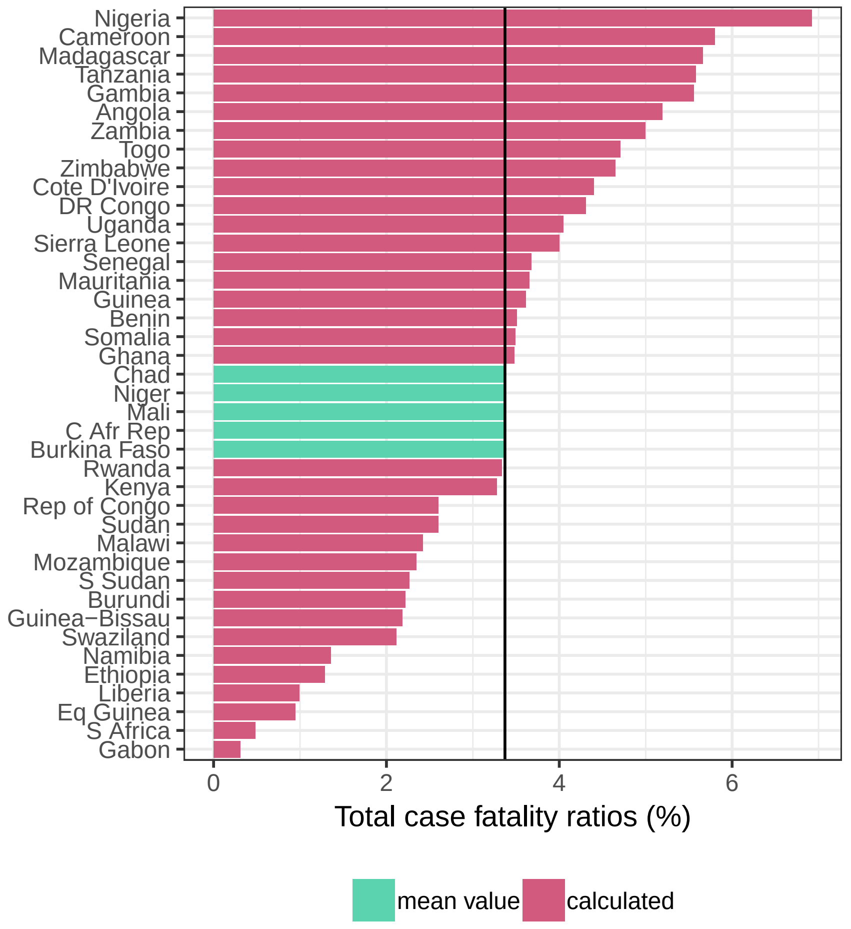

Figure S9: **Case fatality ratios pooled across all available data by country.** We calculated the total case fatality ratios by country for all years of data available in the WHO Global Health Observatory database. For a given country, data could have been reported for any year within the range of 1970 through 2016. For countries with implied case fatality ratios greater than 7% (colored in seafoam green), we used the mean of the case fatality ratios below 7% (3.4%).

We compared the country-level total (pooled across years) CFR estimates to the outputs from country-specific random effects models, where CFR observations were computed annually with all available WHO Global Health Observatory data (Figure S10). We found that the estimates from the random effects models were systematically higher than the total CFRs. We preferred to use the total CFR estimates because they would generate more conservative estimates in the number of deaths averted through vaccination campaigns.

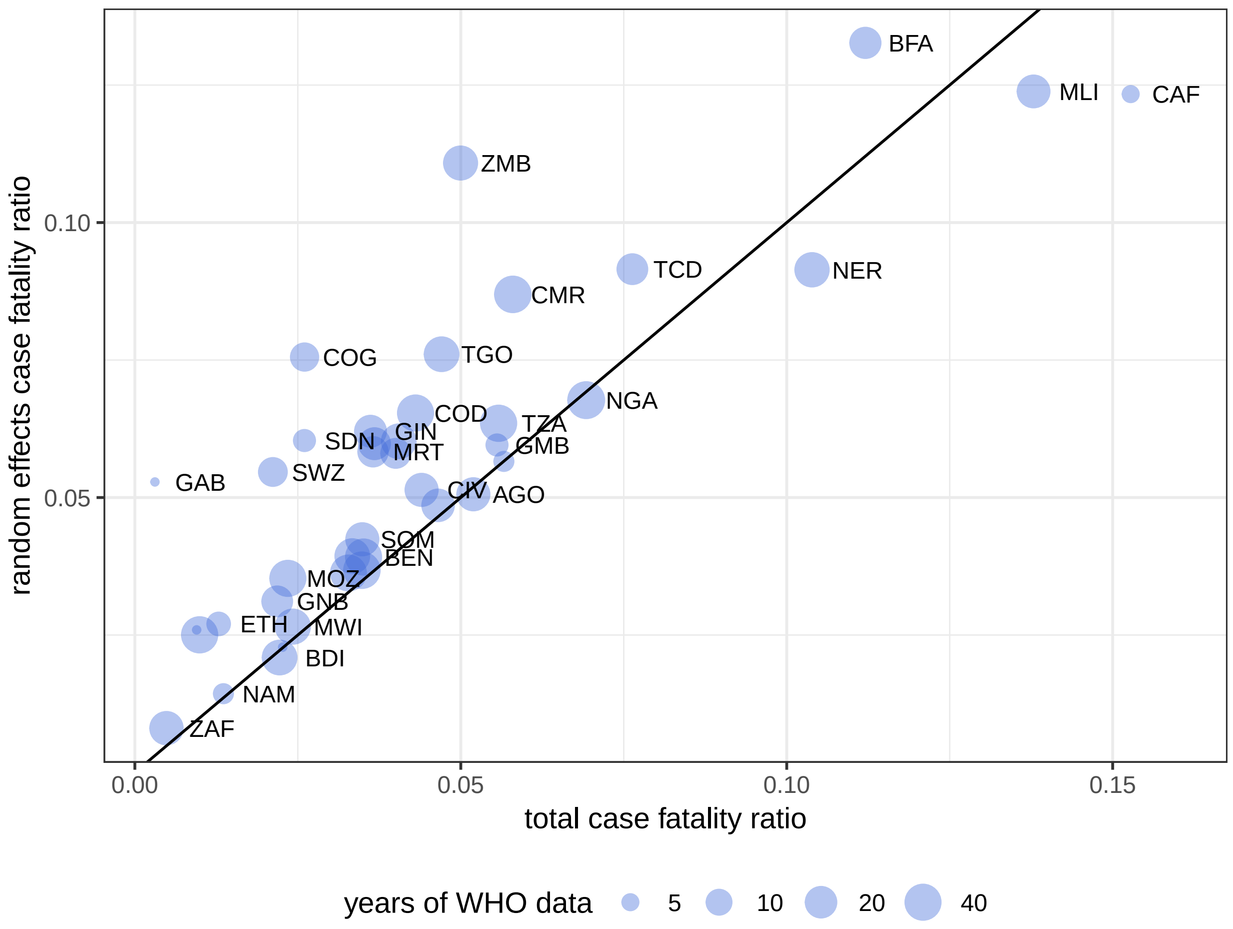

Figure S10: **Comparison of total CFR and random effects model CFR estimates by country.** Random effects model estimates were systematically larger than total CFR estimates, and they were specifically higher in several countries where rate-logistics optimized targeting takes place, such as DR Congo (COD), Cameroon (CMR), and Sierra Leone (SLE).

### Vaccination Campaign Costs

We reviewed four cost surveys for mass OCV campaigns that reported the vaccine delivery and vaccine procurement costs per fully vaccinated person (i.e., receiving two vaccine doses) in non-refugee African settings (Ilboudo and Le Gargasson 2017; Ciglenecki et al. 2013; Cavailler et al. 2006; Schaetti et al. 2012) (Table S2). We excluded the estimate from Schaetti et al. 2012, due to its excessively high vaccine purchase prices. We felt that the exclusion of this study was reasonable based on current policy discussions, which suggest that future, preventive use of OCV in these settings are likely to be made available lower, negotiated rates.

All costs were adjusted to 2017 US dollars (USD) according to the World Bank Consumer Price Index. We used the mean of the three remaining cost survey data points ($6.32 per FVP), to calculate the cost per DALY averted results.

Table S2: **Cost survey estimates (adjusted to 2017 USD) per fully vaccinated person (FVP) for previous vaccination campaigns in Africa.**

| Study | Country | Procurement | Delivery | Total |
|---|---|---|---|---|
| Ilboudo and Le Gargasson 2017 | Malawi | 5.32 | 1.98 | 7.30 |

| Study | Country | Procurement | Delivery | Total |
|---|---|---|---|---|
| Ciglenecki et al. 2013 | Guinea | 5.66 | 2.41 | 8.07 |
| Cavailler et al. 2006 | Mozambique | 1.33 | 2.26 | 3.59 |

### Constant Incidence Assumption

Our model assumed that baseline cholera incidence risk remained constant in our projections and we examined the reasonableness of this assumption using data from country-level annual cholera reports to WHO. First, we gathered all available country-level cholera case data from WHO annual cholera reports, which are reported annually in issues of the Weekly Epidemiological Record, for African countries in our study. Cholera case data were available from 1970 through 2017, but data are not complete for all countries and years.

We also gathered country-level population estimates from the 2019 United Nations World Population Projections, which were available in five-year intervals from 1950 through 2020. We fit a log-linear model to data for each country in order to obtain annual country-level estimates, and then we divided the case and population data to obtain annual estimates of the cholera incidence rate by country and year.

For each country-year pair from 2008-2017, we then compared the observed incidence rate to the expected incidence rate, which was calculated as the country's mean annual incidence rate across all remaining years in 10-year range (Figure S11). The mean of the difference between observed and expected incidence rates (i.e., bias), was nearly zero (1.5E-16); this means that the mean annual incidence was an unbiased predictor of incidence in the held-out year, suggesting that our analysis is valid in the expectation. In addition, there was a positive relationship between the expected and observed incidence rates ($H_0 : \rho = 0$; $\rho = 0.31$, p-value $= 2.4E - 6$).

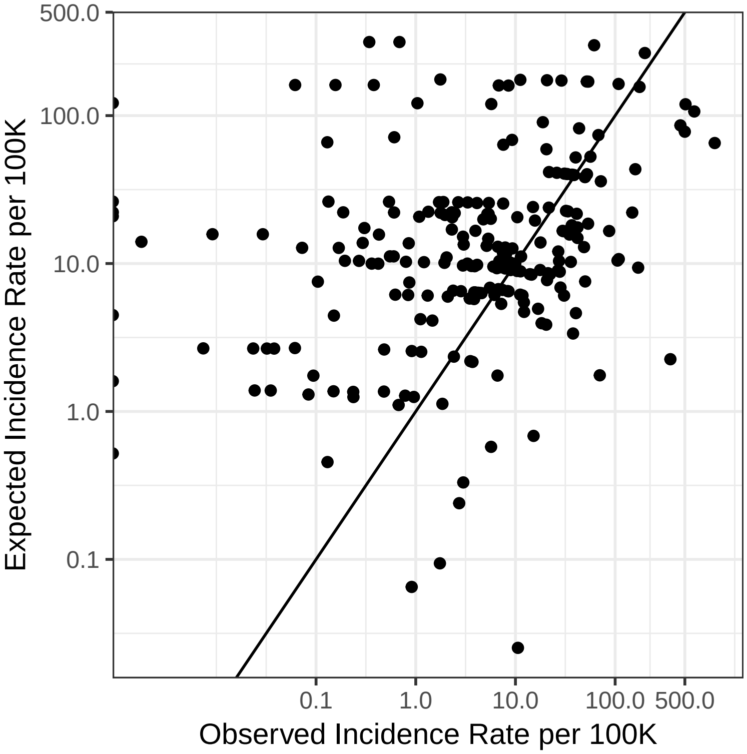

Figure S11: **Observed versus expected mean annual incidence for country-year pairs according to WHO Annual Cholera Reports, 2008-2017.**

### Percentage of Targets in Epidemic Countries

We examined the percentage of targets in the rate optimized, rate-logistics optimized, water optimized, and sanitation optimized strategies that occurred in countries with a coefficient of variation greater than 1.5 in their annual incidence rate from 2008 to 2017. We expect that countries with large coefficients of variation experience “epidemic” (instead of “endemic”) cholera epidemiology, according to a framework previously proposed in Figure 4 of Lessler and Moore et al. (2018).

Table S3: **Percentage of targets in epidemic countries for baseline parameters by vaccination deployment strategy.** Epidemic countries are defined as those with a coefficient of variation in annual incidence greater than 1.5 from 2008-2017, according to cholera data from the WHO annual cholera reports and population data from the UN World Population Prospects.

| Deployment Strategy | Total Targets | # Epidemic Targets | % Epidemic Targets |
|---|---|---|---|
| case-logistics optimized | 225 | 65 | 28.9 |
| rate-logistics optimized | 217 | 59 | 27.2 |
| case optimized | 236 | 69 | 29.2 |
| rate optimized | 227 | 67 | 29.5 |
| sanitation optimized | 253 | 40 | 15.8 |
| watsan optimized | 273 | 4 | 1.5 |
| water optimized | 333 | 19 | 5.7 |

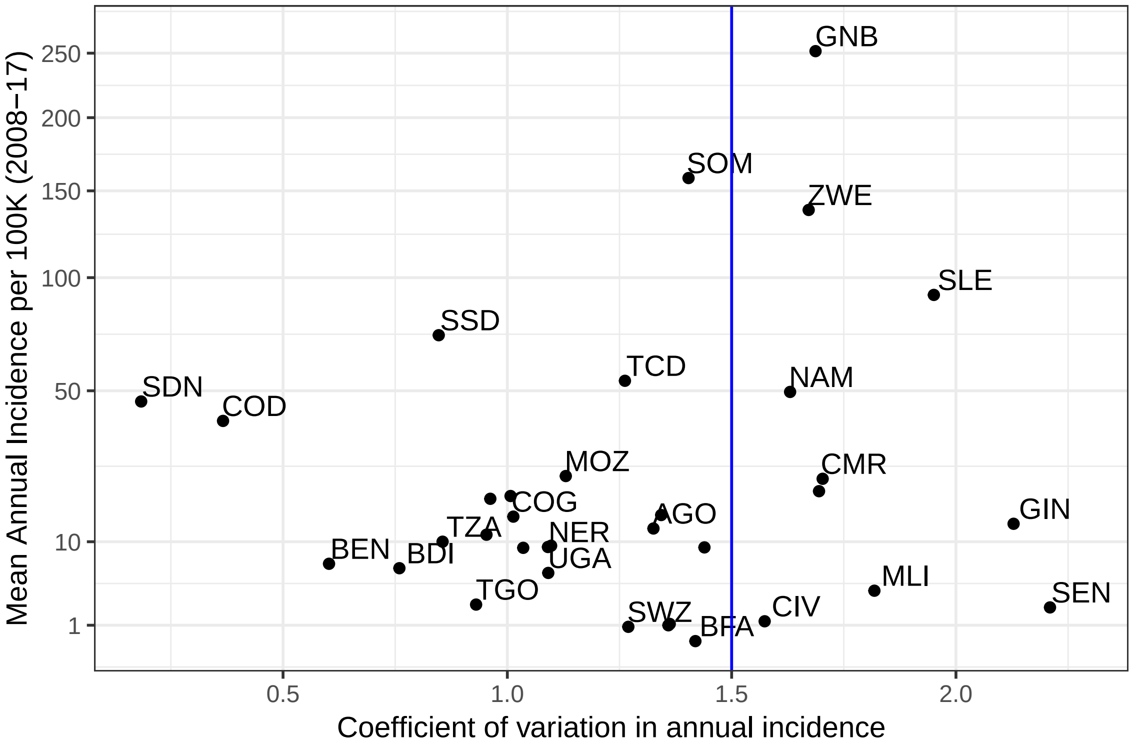

Figure S12: **Cholera incidence versus the coefficient of variation of the annual incidence.** Mean annual incidence per 100,000 people versus the coefficient of variation of the annual incidence from 2008 to 2017 for 35 African countries based on reports to WHO. Countries with one or fewer years of reported data in this period were excluded. The blue line represents the threshold used to delineate endemic (left) and epidemic (right) countries.

In both the rate optimized and rate-logistics optimized strategies, 26% of locations receiving vaccine were located in countries with epidemic dynamics, respectively. In the water optimized and sanitation optimized strategies, 6% and 16% of locations receiving vaccine were located in countries with epidemic dynamics, respectively. These analyses appear in the Rmd supplement file under Percentage of Targets in Epidemic Countries.

### Baseline Model Outcomes

This section presents cumulative cases averted, deaths averted, and DALYs averted for the baseline parameter models for all vaccination deployment scenarios (Figure S13, Figure S14, Figure S15).

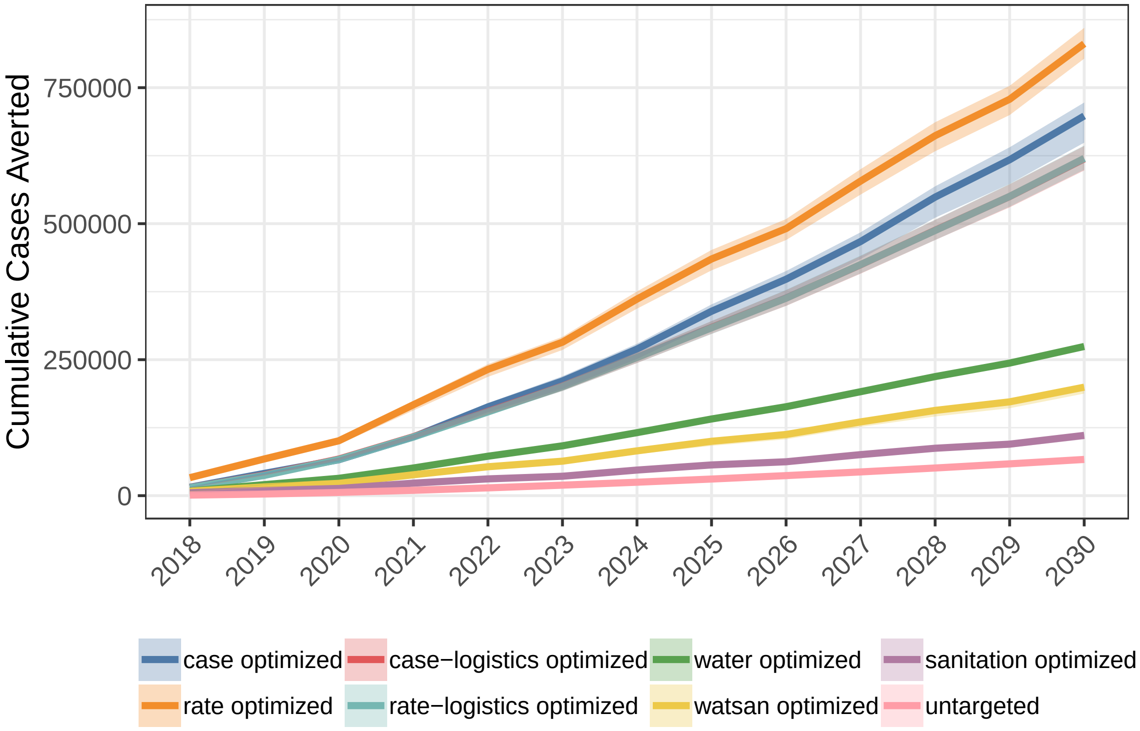

Figure S13: **Cumulative cases averted from mass oral cholera vaccination campaigns across all vaccination deployment strategies in sub-Saharan Africa from 2018 through 2030.**

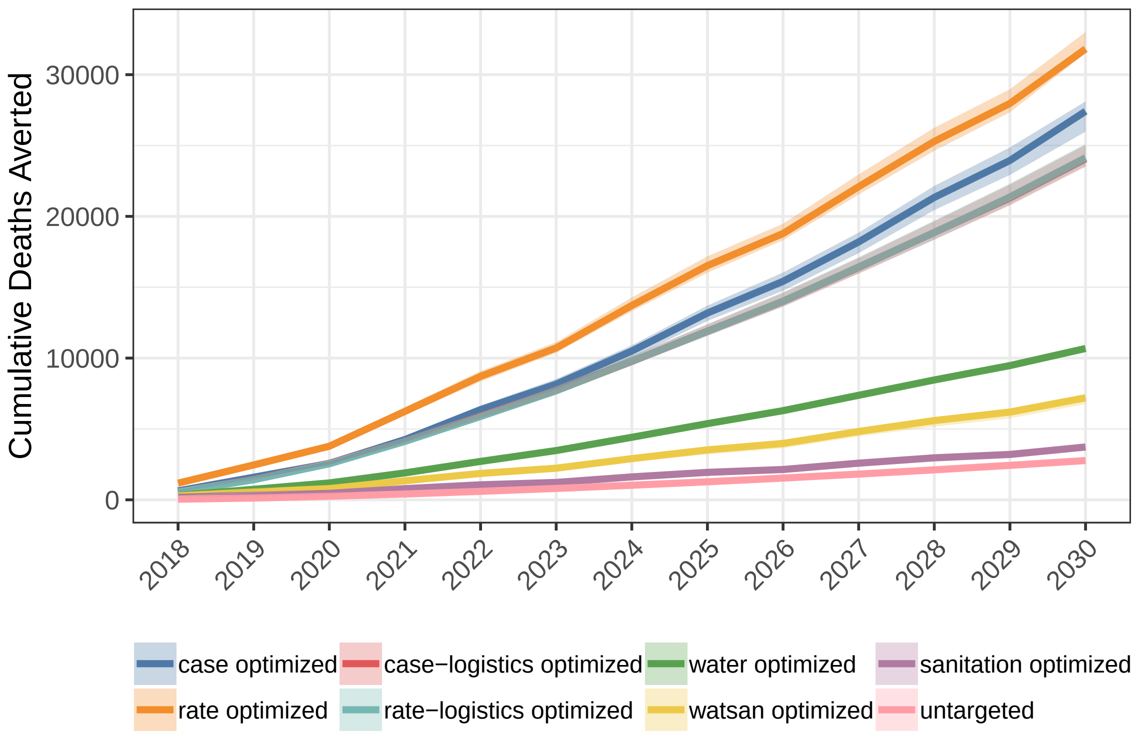

Figure S14: **Cumulative deaths averted from mass oral cholera vaccination campaigns across all vaccination deployment strategies in sub-Saharan Africa from 2018 through 2030.**

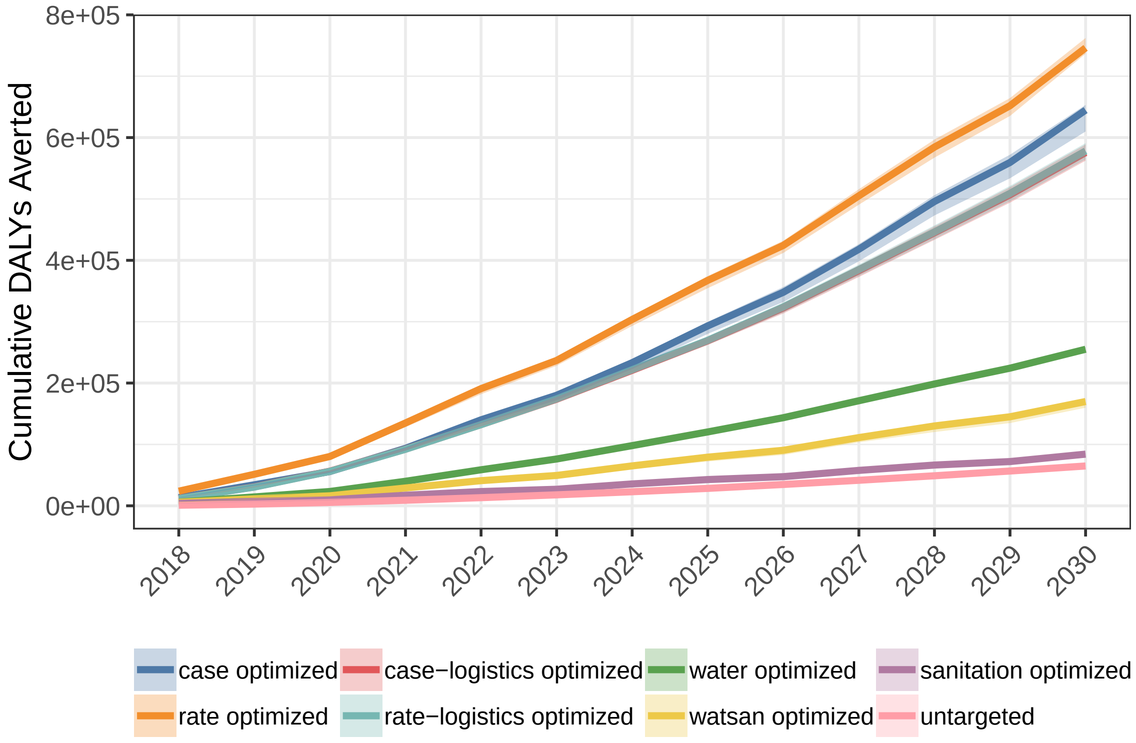

Figure S15: **Cumulative DALYs averted from mass oral cholera vaccination campaigns across all vaccination deployment strategies in sub-Saharan Africa from 2018 through 2030.**

### Summary Tables by Parameter Set

This section presents summary tables of model outcomes by sensitivity parameter set.

#### Baseline

Table S4: **Summary of cumulative health impacts and total cost per DALY averted (USD 2017) from 2018-2030 by vaccination deployment strategy for baseline model parameters.**

| Deployment Strategy | Measure | Mean | 2.5% | 97.5% |
|---|---|---|---|---|
| case-logistics optimized | cases averted | 616188 | 597745 | 642740 |
| case optimized | cases averted | 693007 | 648877 | 722717 |
| rate-logistics optimized | cases averted | 617424 | 599150 | 643791 |
| rate optimized | cases averted | 828971 | 803370 | 859980 |
| sanitation optimized | cases averted | 109817 | 103735 | 114110 |
| untargeted | cases averted | 66562 | 65733 | 67702 |
| water optimized | cases averted | 273939 | 270319 | 277002 |
| watsan optimized | cases averted | 197714 | 187610 | 203934 |
| case-logistics optimized | deaths averted | 24104 | 23501 | 25039 |
| case optimized | deaths averted | 27192 | 25980 | 28116 |
| rate-logistics optimized | deaths averted | 24189 | 23579 | 25115 |
| rate optimized | deaths averted | 31958 | 31503 | 33011 |
| sanitation optimized | deaths averted | 3682 | 3469 | 3812 |
| untargeted | deaths averted | 2771 | 2745 | 2813 |
| water optimized | deaths averted | 10672 | 10517 | 10827 |
| watsan optimized | deaths averted | 7144 | 6760 | 7401 |
| case-logistics optimized | DALYs averted | 575078 | 562373 | 588721 |
| case optimized | DALYs averted | 637975 | 609931 | 653243 |
| rate-logistics optimized | DALYs averted | 577533 | 564572 | 590935 |
| rate optimized | DALYs averted | 746749 | 736607 | 762273 |
| sanitation optimized | DALYs averted | 83228 | 78579 | 86117 |
| untargeted | DALYs averted | 64862 | 64404 | 65532 |
| water optimized | DALYs averted | 255090 | 251723 | 258787 |
| watsan optimized | DALYs averted | 168663 | 159833 | 174676 |
| case-logistics optimized | % cases averted | 25 | 25 | 26 |
| case optimized | % cases averted | 28 | 27 | 29 |
| rate-logistics optimized | % cases averted | 25 | 25 | 26 |
| rate optimized | % cases averted | 34 | 33 | 35 |
| sanitation optimized | % cases averted | 5 | 4 | 5 |
| untargeted | % cases averted | 3 | 3 | 3 |
| water optimized | % cases averted | 11 | 11 | 11 |
| watsan optimized | % cases averted | 8 | 8 | 8 |
| case-logistics optimized | cost per DALY averted | 2393 | 1332 | 3116 |
| case optimized | cost per DALY averted | 2158 | 1199 | 2878 |
| rate-logistics optimized | cost per DALY averted | 2383 | 1327 | 3102 |
| rate optimized | cost per DALY averted | 1843 | 1032 | 2382 |
| sanitation optimized | cost per DALY averted | 16546 | 9121 | 22243 |
| untargeted | cost per DALY averted | 21213 | 11953 | 27263 |
| water optimized | cost per DALY averted | 5394 | 3029 | 6965 |
| watsan optimized | cost per DALY averted | 8164 | 4496 | 10923 |

Table S5: **Mean percentage of cases averted annually from 2018-2030 by vaccination deployment strategy for baseline model parameters.**

| Deployment Strategy | 2018 | 2019 | 2020 | 2021 | 2022 | 2023 | 2024 | 2025 | 2026 | 2027 | 2028 | 2029 | 2030 |
|---|---|---|---|---|---|---|---|---|---|---|---|---|---|
| case-logistics optimized | 9 | 14 | 17 | 24 | 26 | 25 | 28 | 29 | 27 | 30 | 30 | 29 | 31 |
| case optimized | 10 | 15 | 15 | 24 | 31 | 26 | 31 | 36 | 29 | 34 | 39 | 31 | 37 |
| rate-logistics optimized | 9 | 14 | 17 | 24 | 26 | 26 | 29 | 28 | 28 | 30 | 30 | 29 | 31 |
| rate optimized | 21 | 21 | 20 | 39 | 37 | 27 | 43 | 38 | 28 | 43 | 40 | 31 | 47 |
| sanitation optimized | 3 | 3 | 2 | 6 | 4 | 3 | 6 | 5 | 3 | 7 | 6 | 3 | 7 |
| untargeted | 1 | 1 | 2 | 2 | 3 | 3 | 3 | 3 | 3 | 3 | 3 | 4 | 4 |
| water optimized | 6 | 7 | 7 | 11 | 12 | 11 | 13 | 13 | 11 | 14 | 13 | 12 | 14 |
| watsan optimized | 5 | 5 | 4 | 9 | 8 | 6 | 10 | 9 | 6 | 11 | 10 | 7 | 12 |

**Vaccine Efficacy**

Table S6: **Summary of cumulative health impacts and total cost per DALY averted from 2018-2030 by vaccination deployment strategy for low vaccine efficacy model parameters.**

| Deployment Strategy | Measure | Mean | 2.5% | 97.5% |
|---|---|---|---|---|
| case-logistics optimized | cases averted | 326515 | 316982 | 340464 |
| case optimized | cases averted | 369137 | 345440 | 384869 |
| rate-logistics optimized | cases averted | 327201 | 317673 | 341095 |
| rate optimized | cases averted | 438899 | 426065 | 454912 |
| sanitation optimized | cases averted | 58367 | 55054 | 60609 |
| untargeted | cases averted | 44065 | 43516 | 44820 |
| water optimized | cases averted | 144668 | 142815 | 146277 |
| watsan optimized | cases averted | 105076 | 99749 | 108439 |
| case-logistics optimized | deaths averted | 12788 | 12473 | 13283 |
| case optimized | deaths averted | 14486 | 13831 | 14965 |
| rate-logistics optimized | deaths averted | 12836 | 12516 | 13325 |
| rate optimized | deaths averted | 16931 | 16702 | 17470 |
| sanitation optimized | deaths averted | 1948 | 1835 | 2017 |
| untargeted | deaths averted | 1835 | 1817 | 1862 |
| water optimized | deaths averted | 5643 | 5562 | 5724 |
| watsan optimized | deaths averted | 3809 | 3606 | 3947 |
| case-logistics optimized | DALYs averted | 303989 | 297462 | 311136 |
| case optimized | DALYs averted | 339044 | 323911 | 347065 |
| rate-logistics optimized | DALYs averted | 305332 | 298553 | 312372 |
| rate optimized | DALYs averted | 394463 | 389749 | 402247 |
| sanitation optimized | DALYs averted | 44009 | 41524 | 45542 |
| untargeted | DALYs averted | 42856 | 42553 | 43297 |
| water optimized | DALYs averted | 133941 | 132192 | 135859 |
| watsan optimized | DALYs averted | 89666 | 85059 | 92907 |
| case-logistics optimized | % cases averted | 13 | 13 | 14 |
| case optimized | % cases averted | 15 | 14 | 16 |
| rate-logistics optimized | % cases averted | 13 | 13 | 14 |
| rate optimized | % cases averted | 18 | 18 | 18 |
| sanitation optimized | % cases averted | 2 | 2 | 2 |
| untargeted | % cases averted | 2 | 2 | 2 |
| water optimized | % cases averted | 6 | 6 | 6 |
| watsan optimized | % cases averted | 4 | 4 | 4 |
| case-logistics optimized | cost per DALY averted | 4527 | 2521 | 5890 |
| case optimized | cost per DALY averted | 4061 | 2257 | 5419 |
| rate-logistics optimized | cost per DALY averted | 4507 | 2512 | 5864 |
| rate optimized | cost per DALY averted | 3488 | 1955 | 4500 |
| sanitation optimized | cost per DALY averted | 31290 | 17256 | 42083 |
| untargeted | cost per DALY averted | 32106 | 18091 | 41263 |
| water optimized | cost per DALY averted | 10273 | 5771 | 13261 |
| watsan optimized | cost per DALY averted | 15356 | 8451 | 20533 |

Table S7: **Summary of cumulative health impacts and total cost per DALY averted from 2018-2030 by vaccination deployment strategy for high vaccine efficacy model parameters.**

| Deployment Strategy | Measure | Mean | 2.5% | 97.5% |
|---|---|---|---|---|
| case-logistics optimized | cases averted | 776640 | 753350 | 810300 |
| case optimized | cases averted | 874109 | 818049 | 912103 |
| rate-logistics optimized | cases averted | 778189 | 755252 | 811747 |
| rate optimized | cases averted | 1041303 | 1009714 | 1079703 |
| sanitation optimized | cases averted | 137740 | 129874 | 143224 |
| untargeted | cases averted | 87228 | 86142 | 88722 |
| water optimized | cases averted | 344465 | 339953 | 348321 |
| watsan optimized | cases averted | 247919 | 235089 | 255750 |
| case-logistics optimized | deaths averted | 30383 | 29628 | 31572 |
| case optimized | deaths averted | 34282 | 32744 | 35456 |
| rate-logistics optimized | deaths averted | 30492 | 29723 | 31670 |
| rate optimized | deaths averted | 40175 | 39628 | 41474 |
| sanitation optimized | deaths averted | 4615 | 4344 | 4779 |
| untargeted | deaths averted | 3632 | 3598 | 3687 |
| water optimized | deaths averted | 13428 | 13233 | 13622 |
| watsan optimized | deaths averted | 8959 | 8471 | 9283 |
| case-logistics optimized | DALYs averted | 724350 | 708430 | 741730 |
| case optimized | DALYs averted | 803911 | 768384 | 823453 |
| rate-logistics optimized | DALYs averted | 727449 | 711170 | 744616 |
| rate optimized | DALYs averted | 938353 | 926143 | 957303 |
| sanitation optimized | DALYs averted | 104299 | 98360 | 107962 |
| untargeted | DALYs averted | 85100 | 84499 | 85979 |
| water optimized | DALYs averted | 320492 | 316223 | 325113 |
| watsan optimized | DALYs averted | 211291 | 200049 | 218887 |
| case-logistics optimized | % cases averted | 32 | 31 | 33 |
| case optimized | % cases averted | 36 | 34 | 37 |
| rate-logistics optimized | % cases averted | 32 | 31 | 33 |
| rate optimized | % cases averted | 43 | 42 | 44 |
| sanitation optimized | % cases averted | 6 | 5 | 6 |
| untargeted | % cases averted | 4 | 4 | 4 |
| water optimized | % cases averted | 14 | 14 | 14 |
| watsan optimized | % cases averted | 10 | 10 | 10 |
| case-logistics optimized | cost per DALY averted | 1900 | 1057 | 2474 |
| case optimized | cost per DALY averted | 1713 | 951 | 2284 |
| rate-logistics optimized | cost per DALY averted | 1892 | 1053 | 2463 |
| rate optimized | cost per DALY averted | 1466 | 821 | 1894 |
| sanitation optimized | cost per DALY averted | 13203 | 7277 | 17774 |
| untargeted | cost per DALY averted | 16169 | 9110 | 20780 |
| water optimized | cost per DALY averted | 4293 | 2412 | 5543 |
| watsan optimized | cost per DALY averted | 6517 | 3588 | 8727 |

**Indirect Vaccine Protection**

Table S8: **Summary of cumulative health impacts and total cost per DALY averted from 2018-2030 by vaccination deployment strategy for a model with no indirect effects.**

| Deployment Strategy | Measure | Mean | 2.5% | 97.5% |
|---|---|---|---|---|
| case-logistics optimized | cases averted | 393974 | 382279 | 410984 |
| case optimized | cases averted | 444490 | 416182 | 463785 |
| rate-logistics optimized | cases averted | 394674 | 383061 | 411605 |
| rate optimized | cases averted | 529020 | 512701 | 548680 |
| sanitation optimized | cases averted | 70144 | 66157 | 72894 |
| untargeted | cases averted | 66066 | 65243 | 67197 |
| water optimized | cases averted | 174812 | 172542 | 176762 |
| watsan optimized | cases averted | 126317 | 119763 | 130363 |
| case-logistics optimized | deaths averted | 15414 | 15026 | 16017 |
| case optimized | deaths averted | 17433 | 16659 | 18032 |
| rate-logistics optimized | deaths averted | 15466 | 15074 | 16062 |
| rate optimized | deaths averted | 20399 | 20106 | 21066 |
| sanitation optimized | deaths averted | 2347 | 2210 | 2430 |
| untargeted | deaths averted | 2751 | 2725 | 2792 |
| water optimized | deaths averted | 6813 | 6715 | 6911 |
| watsan optimized | deaths averted | 4569 | 4319 | 4736 |
| case-logistics optimized | DALYs averted | 367147 | 359046 | 375900 |
| case optimized | DALYs averted | 408695 | 390801 | 418639 |
| rate-logistics optimized | DALYs averted | 368626 | 360381 | 377295 |
| rate optimized | DALYs averted | 476104 | 469564 | 485864 |
| sanitation optimized | DALYs averted | 53035 | 50036 | 54897 |
| untargeted | DALYs averted | 64381 | 63927 | 65046 |
| water optimized | DALYs averted | 162335 | 160188 | 164661 |
| watsan optimized | DALYs averted | 107688 | 101966 | 111589 |
| case-logistics optimized | % cases averted | 16 | 16 | 17 |
| case optimized | % cases averted | 18 | 17 | 19 |
| rate-logistics optimized | % cases averted | 16 | 16 | 17 |
| rate optimized | % cases averted | 22 | 21 | 22 |
| sanitation optimized | % cases averted | 3 | 3 | 3 |
| untargeted | % cases averted | 3 | 3 | 3 |
| water optimized | % cases averted | 7 | 7 | 7 |
| watsan optimized | % cases averted | 5 | 5 | 5 |
| case-logistics optimized | cost per DALY averted | 3748 | 2086 | 4880 |
| case optimized | cost per DALY averted | 3369 | 1871 | 4492 |
| rate-logistics optimized | cost per DALY averted | 3733 | 2079 | 4859 |
| rate optimized | cost per DALY averted | 2890 | 1618 | 3736 |
| sanitation optimized | cost per DALY averted | 25966 | 14315 | 34927 |
| untargeted | cost per DALY averted | 21372 | 12042 | 27467 |
| water optimized | cost per DALY averted | 8476 | 4762 | 10943 |
| watsan optimized | cost per DALY averted | 12786 | 7038 | 17125 |

Table S9: **Summary of cumulative health impacts and total cost per DALY averted from 2018-2030 by vaccination deployment strategy for high indirect effects model parameters.**

| Deployment Strategy | Measure | Mean | 2.5% | 97.5% |
|---|---|---|---|---|
| case-logistics optimized | cases averted | 851603 | 826476 | 888509 |
| case optimized | cases averted | 963346 | 901803 | 1005654 |
| rate-logistics optimized | cases averted | 853018 | 828029 | 889882 |
| rate optimized | cases averted | 1141545 | 1106561 | 1183582 |
| sanitation optimized | cases averted | 151537 | 142706 | 157630 |
| untargeted | cases averted | 630214 | 622362 | 641010 |
| water optimized | cases averted | 377128 | 372274 | 381323 |
| watsan optimized | cases averted | 272743 | 258422 | 281626 |
| case-logistics optimized | deaths averted | 33324 | 32487 | 34639 |
| case optimized | deaths averted | 37770 | 36091 | 39081 |
| rate-logistics optimized | deaths averted | 33436 | 32589 | 34735 |
| rate optimized | deaths averted | 44032 | 43408 | 45455 |
| sanitation optimized | deaths averted | 5063 | 4765 | 5243 |
| untargeted | deaths averted | 26239 | 25993 | 26637 |
| water optimized | deaths averted | 14706 | 14495 | 14914 |
| watsan optimized | deaths averted | 9872 | 9326 | 10236 |
| case-logistics optimized | DALYs averted | 792707 | 775300 | 811740 |
| case optimized | DALYs averted | 884863 | 846119 | 906590 |
| rate-logistics optimized | DALYs averted | 795833 | 777984 | 814787 |
| rate optimized | DALYs averted | 1026750 | 1012775 | 1047432 |
| sanitation optimized | DALYs averted | 114361 | 107816 | 118411 |
| untargeted | DALYs averted | 608387 | 604096 | 614560 |
| water optimized | DALYs averted | 349479 | 344863 | 354419 |
| watsan optimized | DALYs averted | 232364 | 219850 | 240889 |
| case-logistics optimized | % cases averted | 35 | 34 | 36 |
| case optimized | % cases averted | 40 | 37 | 41 |
| rate-logistics optimized | % cases averted | 35 | 34 | 36 |
| rate optimized | % cases averted | 47 | 46 | 48 |
| sanitation optimized | % cases averted | 6 | 6 | 6 |
| untargeted | % cases averted | 26 | 26 | 26 |
| water optimized | % cases averted | 15 | 15 | 16 |
| watsan optimized | % cases averted | 11 | 11 | 11 |
| case-logistics optimized | cost per DALY averted | 1736 | 966 | 2260 |
| case optimized | cost per DALY averted | 1556 | 864 | 2075 |
| rate-logistics optimized | cost per DALY averted | 1729 | 963 | 2250 |
| rate optimized | cost per DALY averted | 1340 | 750 | 1732 |
| sanitation optimized | cost per DALY averted | 12042 | 6637 | 16209 |
| untargeted | cost per DALY averted | 2262 | 1275 | 2907 |
| water optimized | cost per DALY averted | 3937 | 2212 | 5082 |
| watsan optimized | cost per DALY averted | 5926 | 3260 | 7943 |

**Campaign Frequency**

Table S10: **Summary of cumulative health impacts and total cost per DALY averted from 2018-2030 by vaccination deployment strategy for the low campaign frequency (every 5 years) parameter set.**

| Deployment Strategy | Measure | Mean | 2.5% | 97.5% |
|---|---|---|---|---|
| case-logistics optimized | cases averted | 414603 | 399415 | 425447 |
| case optimized | cases averted | 499738 | 491281 | 509616 |
| rate-logistics optimized | cases averted | 414705 | 399255 | 425104 |
| rate optimized | cases averted | 568645 | 561335 | 578023 |
| sanitation optimized | cases averted | 88174 | 83557 | 91277 |
| untargeted | cases averted | 66562 | 65733 | 67702 |
| water optimized | cases averted | 197050 | 195011 | 200476 |
| watsan optimized | cases averted | 142130 | 136703 | 146000 |
| case-logistics optimized | deaths averted | 16553 | 15891 | 17589 |
| case optimized | deaths averted | 20134 | 19256 | 21169 |
| rate-logistics optimized | deaths averted | 16566 | 15885 | 17574 |
| rate optimized | deaths averted | 22703 | 21706 | 23625 |
| sanitation optimized | deaths averted | 2993 | 2838 | 3104 |
| untargeted | deaths averted | 2771 | 2745 | 2813 |
| water optimized | deaths averted | 7740 | 7654 | 7872 |
| watsan optimized | deaths averted | 5153 | 4948 | 5307 |
| case-logistics optimized | DALYs averted | 388991 | 375058 | 410773 |
| case optimized | DALYs averted | 469219 | 450203 | 491873 |
| rate-logistics optimized | DALYs averted | 389509 | 375104 | 410722 |
| rate optimized | DALYs averted | 528588 | 510182 | 548806 |
| sanitation optimized | DALYs averted | 70269 | 66677 | 72882 |
| untargeted | DALYs averted | 64862 | 64404 | 65532 |
| water optimized | DALYs averted | 183303 | 181155 | 186502 |
| watsan optimized | DALYs averted | 121593 | 117068 | 125017 |
| case-logistics optimized | % cases averted | 17 | 17 | 18 |
| case optimized | % cases averted | 21 | 20 | 21 |
| rate-logistics optimized | % cases averted | 17 | 17 | 18 |
| rate optimized | % cases averted | 23 | 23 | 24 |
| sanitation optimized | % cases averted | 4 | 3 | 4 |
| untargeted | % cases averted | 3 | 3 | 3 |
| water optimized | % cases averted | 8 | 8 | 8 |
| watsan optimized | % cases averted | 6 | 6 | 6 |
| case-logistics optimized | cost per DALY averted | 3540 | 1906 | 4668 |
| case optimized | cost per DALY averted | 2935 | 1593 | 3880 |
| rate-logistics optimized | cost per DALY averted | 3536 | 1906 | 4665 |
| rate optimized | cost per DALY averted | 2605 | 1427 | 3438 |
| sanitation optimized | cost per DALY averted | 19596 | 10780 | 26247 |
| untargeted | cost per DALY averted | 21213 | 11953 | 27263 |
| water optimized | cost per DALY averted | 7507 | 4207 | 9679 |
| watsan optimized | cost per DALY averted | 11320 | 6279 | 14954 |

Table S11: **Summary of cumulative health impacts and total cost per DALY averted from 2018-2030 by vaccination deployment strategy for the high campaign frequency (every 2 years) parameter set.**

| Deployment Strategy | Measure | Mean | 2.5% | 97.5% |
|---|---|---|---|---|
| case-logistics optimized | cases averted | 733941 | 706351 | 768438 |
| case optimized | cases averted | 746159 | 705970 | 785180 |
| rate-logistics optimized | cases averted | 733837 | 706125 | 768390 |
| rate optimized | cases averted | 1045568 | 992102 | 1093351 |
| sanitation optimized | cases averted | 133530 | 124489 | 139523 |
| untargeted | cases averted | 66562 | 65733 | 67702 |
| water optimized | cases averted | 328983 | 322016 | 335705 |
| watsan optimized | cases averted | 228621 | 212604 | 238137 |
| case-logistics optimized | deaths averted | 28559 | 27680 | 29796 |
| case optimized | deaths averted | 29064 | 27866 | 30523 |
| rate-logistics optimized | deaths averted | 28573 | 27689 | 29815 |
| rate optimized | deaths averted | 39713 | 38256 | 41442 |
| sanitation optimized | deaths averted | 4576 | 4264 | 4784 |
| untargeted | deaths averted | 2771 | 2745 | 2813 |
| water optimized | deaths averted | 12323 | 12045 | 12584 |
| watsan optimized | deaths averted | 7992 | 7454 | 8297 |
| case-logistics optimized | DALYs averted | 687060 | 666820 | 706338 |
| case optimized | DALYs averted | 679724 | 653678 | 701939 |
| rate-logistics optimized | DALYs averted | 687962 | 667609 | 707329 |
| rate optimized | DALYs averted | 924602 | 890844 | 951704 |
| sanitation optimized | DALYs averted | 102401 | 95531 | 106982 |
| untargeted | DALYs averted | 64862 | 64404 | 65532 |
| water optimized | DALYs averted | 294641 | 288776 | 299894 |
| watsan optimized | DALYs averted | 185382 | 173188 | 192427 |
| case-logistics optimized | % cases averted | 30 | 29 | 31 |
| case optimized | % cases averted | 31 | 29 | 32 |
| rate-logistics optimized | % cases averted | 30 | 29 | 31 |
| rate optimized | % cases averted | 43 | 41 | 44 |
| sanitation optimized | % cases averted | 5 | 5 | 6 |
| untargeted | % cases averted | 3 | 3 | 3 |
| water optimized | % cases averted | 14 | 13 | 14 |
| watsan optimized | % cases averted | 9 | 9 | 10 |
| case-logistics optimized | cost per DALY averted | 2003 | 1111 | 2629 |
| case optimized | cost per DALY averted | 2025 | 1119 | 2681 |
| rate-logistics optimized | cost per DALY averted | 2001 | 1109 | 2626 |
| rate optimized | cost per DALY averted | 1489 | 826 | 1969 |
| sanitation optimized | cost per DALY averted | 13453 | 7356 | 18280 |
| untargeted | cost per DALY averted | 21213 | 11953 | 27263 |
| water optimized | cost per DALY averted | 4670 | 2611 | 6066 |
| watsan optimized | cost per DALY averted | 7429 | 4074 | 10072 |

**Campaign Coverage**

Table S12: **Summary of cumulative health impacts and total cost per DALY averted from 2018-2030 by vaccination deployment strategy for low vaccination coverage model parameters.**

| Deployment Strategy | Measure | Mean | 2.5% | 97.5% |
|---|---|---|---|---|
| case-logistics optimized | cases averted | 418326 | 407178 | 431672 |
| case optimized | cases averted | 522825 | 500475 | 535522 |
| rate-logistics optimized | cases averted | 419621 | 408671 | 433002 |
| rate optimized | cases averted | 583203 | 576147 | 592731 |
| sanitation optimized | cases averted | 83844 | 78774 | 86877 |
| untargeted | cases averted | 66562 | 65733 | 67702 |
| water optimized | cases averted | 191337 | 189535 | 193777 |
| watsan optimized | cases averted | 142673 | 138285 | 147706 |
| case-logistics optimized | deaths averted | 16640 | 16227 | 17033 |
| case optimized | deaths averted | 20916 | 20272 | 22079 |
| rate-logistics optimized | deaths averted | 16683 | 16273 | 17075 |
| rate optimized | deaths averted | 23100 | 22143 | 23884 |
| sanitation optimized | deaths averted | 2738 | 2552 | 2843 |
| untargeted | deaths averted | 2771 | 2745 | 2813 |
| water optimized | deaths averted | 7516 | 7433 | 7613 |
| watsan optimized | deaths averted | 5267 | 5069 | 5456 |
| case-logistics optimized | DALYs averted | 394412 | 384971 | 401189 |
| case optimized | DALYs averted | 489638 | 476652 | 515578 |
| rate-logistics optimized | DALYs averted | 395410 | 386278 | 402003 |
| rate optimized | DALYs averted | 540918 | 523208 | 558450 |
| sanitation optimized | DALYs averted | 63536 | 59138 | 65959 |
| untargeted | DALYs averted | 64862 | 64404 | 65532 |
| water optimized | DALYs averted | 179331 | 177509 | 181684 |
| watsan optimized | DALYs averted | 124614 | 120135 | 128890 |
| case-logistics optimized | % cases averted | 17 | 17 | 17 |
| case optimized | % cases averted | 21 | 21 | 22 |
| rate-logistics optimized | % cases averted | 17 | 17 | 18 |
| rate optimized | % cases averted | 24 | 24 | 24 |
| sanitation optimized | % cases averted | 3 | 3 | 4 |
| untargeted | % cases averted | 3 | 3 | 3 |
| water optimized | % cases averted | 8 | 8 | 8 |
| watsan optimized | % cases averted | 6 | 6 | 6 |
| case-logistics optimized | cost per DALY averted | 7757 | 1952 | 20935 |
| case optimized | cost per DALY averted | 6252 | 1522 | 16940 |
| rate-logistics optimized | cost per DALY averted | 7768 | 1947 | 20881 |
| rate optimized | cost per DALY averted | 5674 | 1405 | 15476 |
| sanitation optimized | cost per DALY averted | 48636 | 11924 | 136337 |
| untargeted | cost per DALY averted | 47196 | 11962 | 125964 |
| water optimized | cost per DALY averted | 17006 | 4316 | 45615 |
| watsan optimized | cost per DALY averted | 24453 | 6102 | 67274 |

Table S13: **Summary of cumulative health impacts and total cost per DALY averted from 2018-2030 by vaccination deployment strategy for high vaccination coverage model parameters.**

| Deployment Strategy | Measure | Mean | 2.5% | 97.5% |
|---|---|---|---|---|
| case-logistics optimized | cases averted | 691262 | 666546 | 723443 |
| case optimized | cases averted | 732105 | 688897 | 768931 |
| rate-logistics optimized | cases averted | 698111 | 673333 | 730590 |
| rate optimized | cases averted | 953179 | 911923 | 994805 |
| sanitation optimized | cases averted | 119392 | 112105 | 124480 |
| untargeted | cases averted | 66562 | 65733 | 67702 |
| water optimized | cases averted | 309335 | 303988 | 313822 |
| watsan optimized | cases averted | 208116 | 195495 | 214804 |
| case-logistics optimized | deaths averted | 26869 | 26114 | 27985 |
| case optimized | deaths averted | 28568 | 27360 | 29859 |
| rate-logistics optimized | deaths averted | 27177 | 26406 | 28321 |
| rate optimized | deaths averted | 36479 | 35418 | 37975 |
| sanitation optimized | deaths averted | 4102 | 3855 | 4279 |
| untargeted | deaths averted | 2771 | 2745 | 2813 |
| water optimized | deaths averted | 11796 | 11568 | 12013 |
| watsan optimized | deaths averted | 7332 | 6865 | 7574 |
| case-logistics optimized | DALYs averted | 644035 | 626957 | 661557 |
| case optimized | DALYs averted | 669097 | 641934 | 689465 |
| rate-logistics optimized | DALYs averted | 651998 | 634594 | 670096 |
| rate optimized | DALYs averted | 848872 | 824779 | 872312 |
| sanitation optimized | DALYs averted | 92182 | 86766 | 96086 |
| untargeted | DALYs averted | 64862 | 64404 | 65532 |
| water optimized | DALYs averted | 280290 | 275552 | 284962 |
| watsan optimized | DALYs averted | 170781 | 159894 | 176213 |
| case-logistics optimized | % cases averted | 28 | 27 | 29 |
| case optimized | % cases averted | 30 | 28 | 31 |
| rate-logistics optimized | % cases averted | 29 | 28 | 30 |
| rate optimized | % cases averted | 39 | 38 | 40 |
| sanitation optimized | % cases averted | 5 | 5 | 5 |
| untargeted | % cases averted | 3 | 3 | 3 |
| water optimized | % cases averted | 13 | 12 | 13 |
| watsan optimized | % cases averted | 9 | 8 | 9 |
| case-logistics optimized | cost per DALY averted | 4752 | 1187 | 12914 |
| case optimized | cost per DALY averted | 4575 | 1137 | 12622 |
| rate-logistics optimized | cost per DALY averted | 4712 | 1172 | 12753 |
| rate optimized | cost per DALY averted | 3615 | 904 | 9825 |
| sanitation optimized | cost per DALY averted | 33508 | 8205 | 92816 |
| untargeted | cost per DALY averted | 47196 | 11962 | 125964 |
| water optimized | cost per DALY averted | 10881 | 2749 | 29385 |
| watsan optimized | cost per DALY averted | 17850 | 4455 | 50172 |

**Vaccine Supply**

Table S14: **Summary of cumulative health impacts and total cost per DALY averted from 2018-2030 by vaccination deployment strategy for the low vaccine supply (up to 26 million doses in 2030) parameter set.**

| Deployment Strategy | Measure | Mean | 2.5% | 97.5% |
|---|---|---|---|---|
| case-logistics optimized | cases averted | 467436 | 452372 | 487583 |
| case optimized | cases averted | 475185 | 448519 | 503639 |
| rate-logistics optimized | cases averted | 485513 | 468662 | 505693 |
| rate optimized | cases averted | 710856 | 670679 | 744365 |
| sanitation optimized | cases averted | 89180 | 83697 | 92791 |
| untargeted | cases averted | 44162 | 43613 | 44918 |
| water optimized | cases averted | 224957 | 219843 | 229532 |
| watsan optimized | cases averted | 151121 | 140798 | 157186 |
| case-logistics optimized | deaths averted | 18191 | 17686 | 18926 |
| case optimized | deaths averted | 18496 | 17700 | 19560 |
| rate-logistics optimized | deaths averted | 19043 | 18483 | 19782 |
| rate optimized | deaths averted | 26866 | 25753 | 28116 |
| sanitation optimized | deaths averted | 3053 | 2866 | 3176 |
| untargeted | deaths averted | 1839 | 1822 | 1867 |
| water optimized | deaths averted | 8395 | 8183 | 8594 |
| watsan optimized | deaths averted | 5280 | 4930 | 5478 |
| case-logistics optimized | DALYs averted | 434799 | 423910 | 445718 |
| case optimized | DALYs averted | 426640 | 409677 | 443534 |
| rate-logistics optimized | DALYs averted | 456382 | 443814 | 468410 |
| rate optimized | DALYs averted | 620776 | 595037 | 640358 |
| sanitation optimized | DALYs averted | 67935 | 63852 | 70647 |
| untargeted | DALYs averted | 42634 | 42335 | 43064 |
| water optimized | DALYs averted | 198652 | 194172 | 202740 |
| watsan optimized | DALYs averted | 121434 | 113464 | 125924 |
| case-logistics optimized | % cases averted | 19 | 19 | 20 |
| case optimized | % cases averted | 20 | 19 | 20 |
| rate-logistics optimized | % cases averted | 20 | 19 | 20 |
| rate optimized | % cases averted | 29 | 28 | 30 |
| sanitation optimized | % cases averted | 4 | 3 | 4 |
| untargeted | % cases averted | 2 | 2 | 2 |
| water optimized | % cases averted | 9 | 9 | 9 |
| watsan optimized | % cases averted | 6 | 6 | 6 |
| case-logistics optimized | cost per DALY averted | 2048 | 1137 | 2678 |
| case optimized | cost per DALY averted | 2088 | 1148 | 2766 |
| rate-logistics optimized | cost per DALY averted | 1951 | 1082 | 2557 |
| rate optimized | cost per DALY averted | 1435 | 794 | 1908 |
| sanitation optimized | cost per DALY averted | 13117 | 7209 | 17685 |
| untargeted | cost per DALY averted | 20881 | 11768 | 26836 |
| water optimized | cost per DALY averted | 4482 | 2500 | 5840 |
| watsan optimized | cost per DALY averted | 7338 | 4029 | 9954 |

Table S15: **Summary of cumulative health impacts and total cost per DALY averted from 2018-2030 by vaccination deployment strategy for the high vaccine supply (up to 95 million doses in 2030) parameter set.**

| Deployment Strategy | Measure | Mean | 2.5% | 97.5% |
|---|---|---|---|---|
| case-logistics optimized | cases averted | 683745 | 662012 | 702958 |
| case optimized | cases averted | 827238 | 801111 | 846488 |
| rate-logistics optimized | cases averted | 683541 | 661693 | 702757 |
| rate optimized | cases averted | 951412 | 940803 | 966999 |
| sanitation optimized | cases averted | 143553 | 135622 | 148327 |
| untargeted | cases averted | 92891 | 91733 | 94483 |
| water optimized | cases averted | 334830 | 331615 | 339587 |
| watsan optimized | cases averted | 233431 | 224152 | 240073 |
| case-logistics optimized | deaths averted | 27216 | 26333 | 28244 |
| case optimized | deaths averted | 33204 | 32065 | 35005 |
| rate-logistics optimized | deaths averted | 27195 | 26306 | 28222 |
| rate optimized | deaths averted | 37772 | 36251 | 38978 |
| sanitation optimized | deaths averted | 4815 | 4540 | 4990 |
| untargeted | deaths averted | 3867 | 3831 | 3926 |
| water optimized | deaths averted | 13205 | 13064 | 13400 |
| watsan optimized | deaths averted | 8484 | 8143 | 8749 |
| case-logistics optimized | DALYs averted | 645756 | 626824 | 667274 |
| case optimized | DALYs averted | 779348 | 755177 | 819210 |
| rate-logistics optimized | DALYs averted | 645458 | 626468 | 666987 |
| rate optimized | DALYs averted | 885827 | 857653 | 911128 |
| sanitation optimized | DALYs averted | 113092 | 106699 | 117278 |
| untargeted | DALYs averted | 90913 | 90269 | 91861 |
| water optimized | DALYs averted | 317400 | 313809 | 322136 |
| watsan optimized | DALYs averted | 201756 | 194159 | 207630 |
| case-logistics optimized | % cases averted | 28 | 27 | 29 |
| case optimized | % cases averted | 34 | 33 | 35 |
| rate-logistics optimized | % cases averted | 28 | 27 | 29 |
| rate optimized | % cases averted | 39 | 39 | 40 |
| sanitation optimized | % cases averted | 6 | 6 | 6 |
| untargeted | % cases averted | 4 | 4 | 4 |
| water optimized | % cases averted | 14 | 14 | 14 |
| watsan optimized | % cases averted | 10 | 9 | 10 |
| case-logistics optimized | cost per DALY averted | 3016 | 1659 | 3950 |
| case optimized | cost per DALY averted | 2500 | 1353 | 3279 |
| rate-logistics optimized | cost per DALY averted | 3017 | 1660 | 3952 |
| rate optimized | cost per DALY averted | 2199 | 1216 | 2893 |
| sanitation optimized | cost per DALY averted | 17230 | 9477 | 23202 |
| untargeted | cost per DALY averted | 21414 | 12064 | 27521 |
| water optimized | cost per DALY averted | 6134 | 3442 | 7902 |
| watsan optimized | cost per DALY averted | 9653 | 5346 | 12756 |

**Population Turnover**

Table S16: **Summary of cumulative health impacts and total cost per DALY averted from 2018-2030 by vaccination deployment strategy for the high population turnover rate/low life expectancy (56 years) parameter set.**

| Deployment Strategy | Measure | Mean | 2.5% | 97.5% |
|---|---|---|---|---|
| case-logistics optimized | cases averted | 613686 | 595328 | 640112 |
| case optimized | cases averted | 690216 | 646263 | 719790 |
| rate-logistics optimized | cases averted | 614866 | 596690 | 641100 |
| rate optimized | cases averted | 825662 | 800179 | 856550 |
| sanitation optimized | cases averted | 109391 | 103335 | 113665 |
| untargeted | cases averted | 66347 | 65520 | 67483 |
| water optimized | cases averted | 272826 | 269222 | 275877 |
| watsan optimized | cases averted | 196941 | 186880 | 203137 |
| case-logistics optimized | deaths averted | 24006 | 23406 | 24937 |
| case optimized | deaths averted | 27083 | 25876 | 28002 |
| rate-logistics optimized | deaths averted | 24089 | 23482 | 25009 |
| rate optimized | deaths averted | 31830 | 31378 | 32879 |
| sanitation optimized | deaths averted | 3668 | 3455 | 3797 |
| untargeted | deaths averted | 2762 | 2737 | 2804 |
| water optimized | deaths averted | 10629 | 10474 | 10783 |
| watsan optimized | deaths averted | 7116 | 6734 | 7372 |
| case-logistics optimized | DALYs averted | 572737 | 560091 | 586310 |
| case optimized | DALYs averted | 635405 | 607466 | 650599 |
| rate-logistics optimized | DALYs averted | 575106 | 562227 | 588428 |
| rate optimized | DALYs averted | 743747 | 733673 | 759205 |
| sanitation optimized | DALYs averted | 82903 | 78273 | 85780 |
| untargeted | DALYs averted | 64652 | 64195 | 65319 |
| water optimized | DALYs averted | 254045 | 250689 | 257727 |
| watsan optimized | DALYs averted | 168007 | 159217 | 173997 |
| case-logistics optimized | % cases averted | 25 | 25 | 26 |
| case optimized | % cases averted | 28 | 27 | 29 |
| rate-logistics optimized | % cases averted | 25 | 25 | 26 |
| rate optimized | % cases averted | 34 | 33 | 35 |
| sanitation optimized | % cases averted | 4 | 4 | 5 |
| untargeted | % cases averted | 3 | 3 | 3 |
| water optimized | % cases averted | 11 | 11 | 11 |
| watsan optimized | % cases averted | 8 | 8 | 8 |
| case-logistics optimized | cost per DALY averted | 2403 | 1337 | 3128 |
| case optimized | cost per DALY averted | 2167 | 1204 | 2889 |
| rate-logistics optimized | cost per DALY averted | 2393 | 1333 | 3116 |
| rate optimized | cost per DALY averted | 1850 | 1036 | 2391 |
| sanitation optimized | cost per DALY averted | 16610 | 9157 | 22330 |
| untargeted | cost per DALY averted | 21283 | 11992 | 27352 |
| water optimized | cost per DALY averted | 5416 | 3042 | 6994 |
| watsan optimized | cost per DALY averted | 8195 | 4513 | 10965 |

Table S17: **Summary of cumulative health impacts and total cost per DALY averted from 2018-2030 by vaccination deployment strategy for the low population turnover rate/high life expectancy (70 years) parameter set.**

| Deployment Strategy | Measure | Mean | 2.5% | 97.5% |
|---|---|---|---|---|
| case-logistics optimized | cases averted | 683745 | 662012 | 702958 |
| case optimized | cases averted | 827238 | 801111 | 846488 |
| rate-logistics optimized | cases averted | 683541 | 661693 | 702757 |
| rate optimized | cases averted | 951412 | 940803 | 966999 |
| sanitation optimized | cases averted | 143553 | 135622 | 148327 |
| untargeted | cases averted | 92891 | 91733 | 94483 |
| water optimized | cases averted | 334830 | 331615 | 339587 |
| watsan optimized | cases averted | 233431 | 224152 | 240073 |
| case-logistics optimized | deaths averted | 27216 | 26333 | 28244 |
| case optimized | deaths averted | 33204 | 32065 | 35005 |
| rate-logistics optimized | deaths averted | 27195 | 26306 | 28222 |
| rate optimized | deaths averted | 37772 | 36251 | 38978 |
| sanitation optimized | deaths averted | 4815 | 4540 | 4990 |
| untargeted | deaths averted | 3867 | 3831 | 3926 |
| water optimized | deaths averted | 13205 | 13064 | 13400 |
| watsan optimized | deaths averted | 8484 | 8143 | 8749 |
| case-logistics optimized | DALYs averted | 645756 | 626824 | 667274 |
| case optimized | DALYs averted | 779348 | 755177 | 819210 |
| rate-logistics optimized | DALYs averted | 645458 | 626468 | 666987 |
| rate optimized | DALYs averted | 885827 | 857653 | 911128 |
| sanitation optimized | DALYs averted | 113092 | 106699 | 117278 |
| untargeted | DALYs averted | 90913 | 90269 | 91861 |
| water optimized | DALYs averted | 317400 | 313809 | 322136 |
| watsan optimized | DALYs averted | 201756 | 194159 | 207630 |
| case-logistics optimized | % cases averted | 28 | 27 | 29 |
| case optimized | % cases averted | 34 | 33 | 35 |
| rate-logistics optimized | % cases averted | 28 | 27 | 29 |
| rate optimized | % cases averted | 39 | 39 | 40 |
| sanitation optimized | % cases averted | 6 | 6 | 6 |
| untargeted | % cases averted | 4 | 4 | 4 |
| water optimized | % cases averted | 14 | 14 | 14 |
| watsan optimized | % cases averted | 10 | 9 | 10 |
| case-logistics optimized | cost per DALY averted | 3016 | 1659 | 3950 |
| case optimized | cost per DALY averted | 2500 | 1353 | 3279 |
| rate-logistics optimized | cost per DALY averted | 3017 | 1660 | 3952 |
| rate optimized | cost per DALY averted | 2199 | 1216 | 2893 |
| sanitation optimized | cost per DALY averted | 17230 | 9477 | 23202 |
| untargeted | cost per DALY averted | 21414 | 12064 | 27521 |
| water optimized | cost per DALY averted | 6134 | 3442 | 7902 |
| watsan optimized | cost per DALY averted | 9653 | 5346 | 12756 |
